# Adolescent Astrocyte Dysregulation Impairs Prefrontal Interneuron Maturation and Adult Cognition

**DOI:** 10.64898/2026.04.26.720933

**Authors:** Johanna Furrer, Viktor Beilmann, Anna Cavaccini, Eliane Boesch, Sina M. Schalbetter, Ali Özgür Argunşah, Sarai Fischer, Aurelia Portmann, Olga Krzyzaniak, Felipe Velasquez Moros, Luca Ravotto, Theofanis Karayannis, Bruno Weber, Tina Notter

**Affiliations:** Institute of Pharmacology and Toxicology, University of Zurich, Zurich, Switzerland; Brain Research Institute, University of Zurich, Zurich, Switzerland; Institute of Veterinary Pharmacology and Toxicology, University of Zurich, Zurich, Switzerland; Neuroscience Center Zurich, University and ETH Zurich, Zurich, Switzerland

**Keywords:** Prefrontal cortex, astrocytes, DREADD, adolescence, behavior, PV interneurons

## Abstract

The prefrontal cortex (PFC), a brain region critical for executive and cognitive functions, is characterized by its protracted maturation extending through adolescence until early adulthood. During adolescence, the PFC undergoes substantial rearrangements, creating a window of heightened plasticity allowing experience-dependent refinement of neural networks. While this extended plasticity supports the development of higher-order cognitive functions, it also confers increased vulnerability to environmental and biological perturbations that can disrupt circuit development and contribute to cognitive and behavioral impairments relevant to psychiatric disorders. Astrocytes are central regulators of brain homeostasis and actively participate in developmental processes that shape postnatal brain maturation. Although astrocyte dysfunction has been increasingly linked to psychiatric pathophysiology, it remains unknown whether aberrant astrocyte activity can directly influence PFC development and cognitive maturation. Here, using selective modulation of astrocyte activity during defined developmental windows in the PFC, we show that abnormal astrocyte activity during adolescence induces transient synaptic loss through enhanced microglial phagocytosis, produces long-lasting alterations in fast-spiking parvalbumin (PV) interneurons, and results in persistent deficits in PFC-dependent behaviors. Together, these findings provide causal evidence that disrupted astrocyte function during adolescent PFC maturation can lead to persistent neuronal and cognitive deficits with relevance to major psychiatric disorders.

## Introduction

The prefrontal cortex (PFC) is unique among mammalian brain regions in that its development continues through adolescence into early adulthood, making it the last region to reach full maturity^1–4^. This protracted maturation supports the development of higher-order cognitive functions, including decision-making, planning, goal-directed behavior, and social cognition^5–8^. While extended plasticity allows for experience-dependent refinement of circuits^9,10^, it also confers heightened vulnerability to environmental perturbations^11,12^. In rodents, adverse experiences during adolescence induce long-lasting structural, functional, and cognitive deficits^13–15^, and in humans, structural and functional abnormalities in the PFC are strongly linked to psychiatric disorders such as schizophrenia (SZ) and bipolar disorder (BD), which typically emerge in late adolescence or early adulthood^16–19^. Accordingly, it has been proposed that environmental insults during adolescence disrupt PFC maturation, contributing to persistent structural, functional, and cognitive deficits^20–22^.

Adolescent PFC maturation is marked by refinement of synaptic connections and circuits^23–25^, maturation of fast-spiking (FS) parvalbumin (PV) interneurons^26–31^, and the emergence of gamma-frequency oscillations^32–34^, all of which are critical for mature cognitive function. Glial cells actively contribute to circuit refinement during postnatal development. Microglia prune synapses in an activity-dependent manner^35–38^, and astrocytes regulate synaptic elimination through IP3-dependent Ca^2+^ mobilization^39^, direct phagocytosis via MERTK and MEGF10^40^, and modulation of microglia-mediated pruning^35,41^. While microglia have been shown to influence prefrontal synaptic and cognitive maturation^42,43^, the role of astrocytes in PFC development is not well defined.

Given their critical role in establishing excitatory–inhibitory balance in the PFC, PV interneuron maturation may be fundamental to the functional development of the PFC. Although they acquire fast-spiking properties during pre-adolescence^27,28^, their excitatory drive and overall maturation, reflected by increased PV expression and the development of perineuronal nets, continue throughout adolescence^26^. These changes coincide with the emergence of gamma-frequency oscillations, which are critical for higher-order cognitive processes^32–34^. Disruption of PV interneuron maturation during this sensitive window can result in long-lasting impairments in gamma oscillatory activity and PFC-dependent behaviors^29^. Nevertheless, the mechanisms underlying PV interneuron maturation remain incompletely understood, and astrocytes may contribute, either directly or indirectly, by regulating perineuronal net formation^44^ or influencing synaptic remodeling and modulating excitatory drive. Notably, transcriptomic analyses in postmortem tissue from SZ and BD patients reveal convergent alterations in PV interneurons and astrocytes, with PV-related genes downregulated and astrocytic genes upregulated, highlighting a potential role for astrocyte–interneuron interactions in shaping prefrontal circuits^45^.

Astrocytes are responsive to various environmental risk factors for psychiatric disorders, such as stress^46^ and cannabinoid exposure^47^, and aberrant prefrontal astrocyte function has been implicated in pathophysiology^45,48–53^. However, it remains unknown whether developmental dysregulation of astrocytes can alter PFC maturation and cognition. To address this knowledge gap, we developed an adolescent DREADD-based astrocyte model (ADAM) that enables cell-selective modulation of prefrontal astrocyte activity during defined developmental windows. Using ADAM, we show that transient overactivation of astrocytes during a critical period of adolescence enhances microglia-mediated synaptic pruning and produces long-lasting alterations in fast-spiking PV interneurons and persistent deficits in PFC-dependent behaviors. These findings establish a causal link between aberrant astrocyte activity during adolescent PFC maturation and enduring functional and cognitive deficits, which are associated with psychiatric disorders.

## Results

### Establishing a DREADD-based model for transient overactivation of prefrontal astrocytes during adolescence

To investigate the role of astrocytes in postnatal PFC maturation, we developed ADAM, an adolescent DREADD-based astrocyte model. This approach enables transient, cell-type-specific stimulation of astrocytes during defined windows of adolescence in freely moving mice. The system utilizes the modified muscarinic Gq-protein-coupled receptor hM3DGq, which elevates intracellular Ca^2+^ via IP3 signaling, a pathway critical for astrocytic function^54^, and previously implicated in developmental synapse elimination in the somatosensory cortex^39^. To achieve region- and astrocyte-specific expression, a recombinant adeno-associated virus (rAAV) encoding hM3DGq under the glial fibrillary acidic protein (GFAP) promoter was bilaterally injected into the medial PFC (mPFC, encompassing anterior cingulate and prelimbic subregions) of 21-day-old (postnatal day, P21) C57BL/6 male mice (**Fig. 1A-B**). During adolescence (between P30 and P50), astrocytes were repeatedly stimulated by daily oral administration of clozapine-*N*-oxide (CNO, 1 mg/kg, **Fig. 1D**), while vehicle (VEH)-treated hM3DGq-expressing mice served as controls. CNO or VEH were delivered non-invasively using the micropipette-guided drug administration (MDA) method^55,56^, avoiding potential confounds from stress associated with repeated injections^57^. Following the final administration, animals were left undisturbed until adulthood (>P84), when long-term outcomes were assessed. Postmortem immunohistochemistry confirmed robust and selective astrocytic expression of hM3DGq, with 92% of S100β⁺ astrocytes expressing hM3DGq, and 97% of hM3DGq⁺ cells co-expressing S100β (**Fig. 1C; Supplementary Fig. S1A-B**).

**Figure 1.**
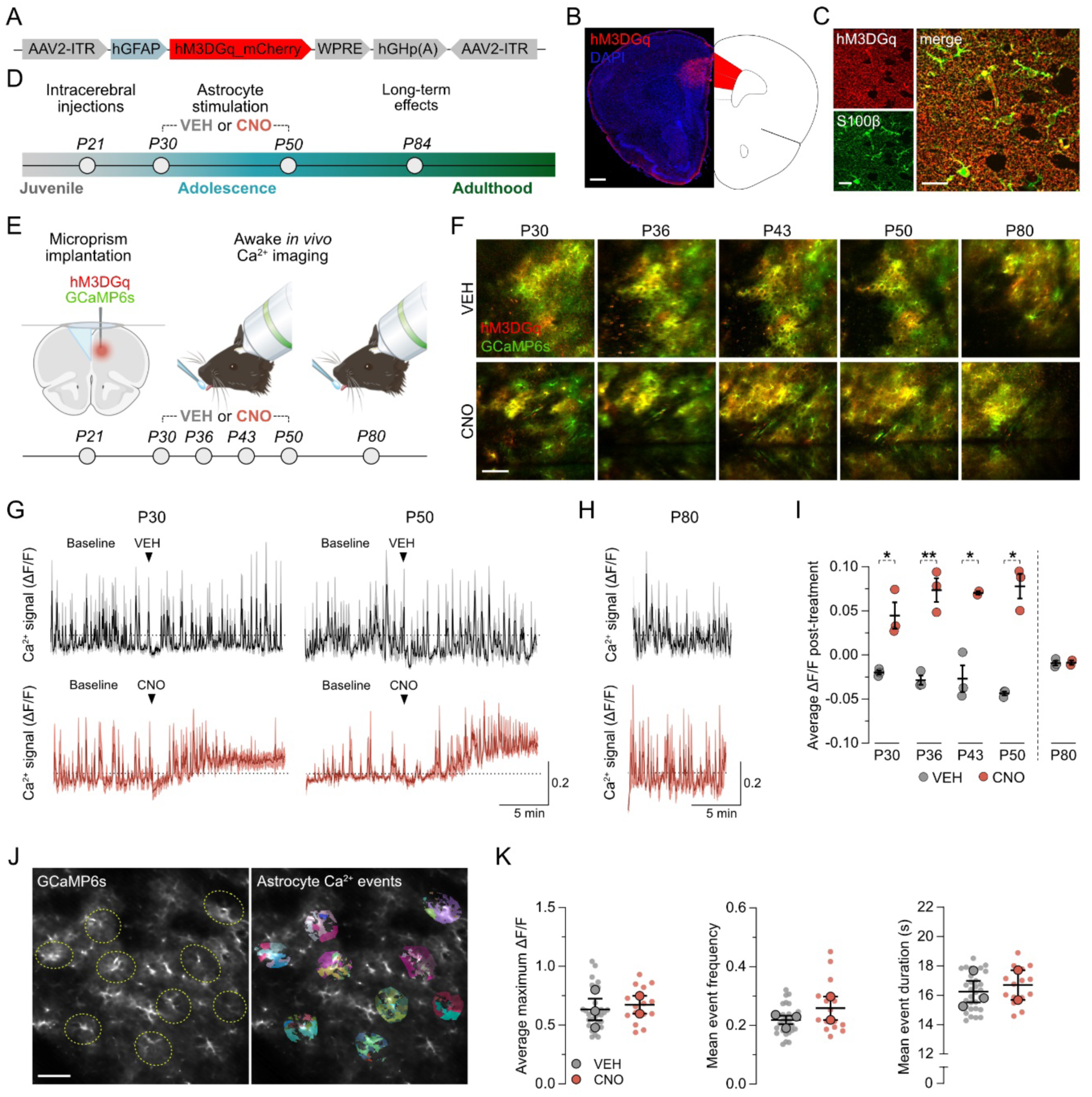
Validation of the adolescent DREADD-based astrocyte model (ADAM) using longitudinal *in vivo* imaging. **(A)** Simplified schematic representation of the recombinant adeno-associated virus (rAAV) expressing hM3DGq-mCherry under the control of the astrocyte-specific hGFAP promoter. **(B)** Representative tile-scan image showing hM3DGq-mCherry expression in adulthood following bilateral stereotaxic rAAV injection into the medial prefrontal cortex (mPFC) at postnatal day (P)21. hM3DGq expression is shown in red; cell nuclei are shown in blue (DAPI). Scale bar = 500 µm. **(C)** Representative double-immunofluorescence images confirming astrocyte-specific expression of hM3DGq (red), as indicated by colocalization with the astrocytic marker S100β (green). Scale bars = 20 µm. **(D)** Experimental timeline of the adolescent DREADD-based astrocyte model (ADAM). Juvenile mice received bilateral stereotaxic injections at P21, followed by daily administration of vehicle (VEH) or clozapine-*N*-oxide (CNO; 1 mg/kg, oral, via MDA) during adolescence (P30-P50). Long-term effects were assessed in adulthood (>P84). **(E)** Schematic overview of the *in vivo* validation of ADAM using two-photon Ca^2+^ imaging in awake mice. At P21, mice underwent microprism implantation and co-injection of rAAVs expressing hM3DGq and the calcium indicator GCaMP6s under the hGFAP promoter. Animals then received daily VEH or CNO from P30 to P50 and were subjected to weekly awake two-photon imaging sessions to monitor hM3DGq-induced astrocytic Ca^2+^ signaling. An additional imaging session was performed in adulthood (P80) to assess potential long-term effects on

### Longitudinal *in vivo* Ca^2+^ imaging confirms pronounced transient astrocyte activation

To verify functional hM3DGq-dependent Ca^2+^ mobilization, we performed longitudinal two-photon imaging in awake mice. A microprism implanted into the longitudinal fissure provided optical access to the mPFC^58^. hM3DGq was co-expressed with the Ca^2+^ indicator GCaMP6s under the GFAP promoter in the hemisphere contralateral to the microprism, allowing simultaneous chemogenetic activation and Ca^2+^ monitoring (**Fig. 1E**). Daily CNO or VEH administration was performed between P30 and P50, with concomitant imaging at P30, P36, P43, and P50 (**Fig. 1E**). A final imaging session at P80 assessed baseline astrocytic activity in adulthood. Successful co-transduction was confirmed at P30 and P80, with 92% of cells co-expressing mCherry and GCaMP6s (**Supplementary Fig. S1C**). Longitudinal imaging enabled repeated measurements of the same astrocytes from adolescence into adulthood (**Fig. 1F**), except for one CNO-treated animal in which imaging at P43 and P80 was not possible due to poor image quality.

During adolescent imaging sessions, each CNO administration produced a robust elevation in astrocytic Ca^2+^ compared to VEH, as reflected by increased mean ΔF/F values (**Fig. 1G-I; Supplementary Fig. S1D**). Importantly, by adulthood, Ca^2+^ dynamics and mean ΔF/F values were indistinguishable between groups (**Fig. 1H,I**). Event-based analysis using AQuA^59^ confirmed the absence of group differences in maximum ΔF/F, event frequency, or duration in adult astrocytes (**Fig. 1J-K**). Consistent with these functional readouts, GFAP immunoreactivity in adulthood was unaltered (**Supplementary Fig. S1E-F**), indicating no evidence of long-term alterations in astrocyte reactivity. Together, these results demonstrate that ADAM enables repeated, temporally controlled stimulation of astrocytes during adolescence without producing lasting alterations in astrocytic Ca^2+^ activity or reactivity in adulthood, providing a robust model system to probe the developmental consequences of transient astrocyte modulation.

### Transient overactivation of prefrontal astrocytes during adolescence leads to long-lasting deficits in PFC-dependent behaviors

We next examined whether transient overactivation of prefrontal astrocytes during adolescence leads to enduring impairments in PFC-dependent behaviors. For this purpose, adult hM3DGq-expressing mice (> P84) that received CNO or VEH from P30-P50 were tested in locomotor, cognitive, and social paradigms (**Fig. 2A**). Open-field testing revealed no differences in total distance traveled or center-zone exploration (**Fig. 2B**), indicating intact locomotor activity and anxiety-like behavior.

**Figure 2.**
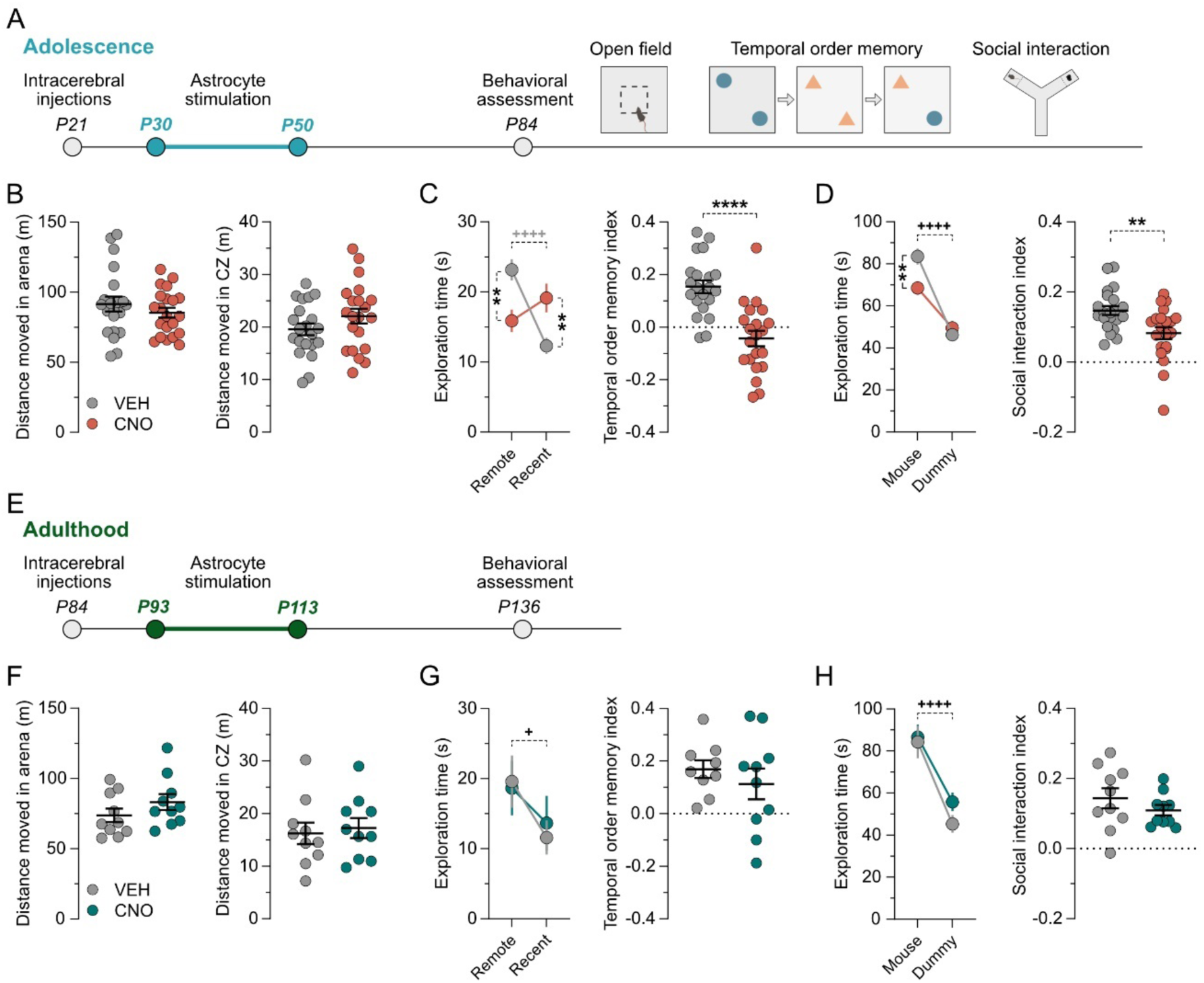
Adolescent prefrontal astrocyte stimulation impairs adult PFC-dependent behaviors. **(A)** Schematic representation of the experimental design used for adolescent manipulations. Animals received bilateral injections at P21, daily VEH or CNO treatment during adolescence from P30 to P50, and were behaviorally assessed in the open field test (OF), the temporal order memory test (TOMT), and the social interaction test (SI) in adulthood (> P84). Panels B-D show the combined results of two independent cohorts (**Supplementary Figure S2**), resulting in a total of *N* = 21 animals per group. **(B)** Total distance moved in the entire arena (left) and center zone (right) during the OF. **(C)** Absolute exploration times of the temporally remote and recent objects (line plots) and temporal order memory index (scatter plot) in the TOMT. Line plot: \*\**p* < 0.01 (reflecting the difference between treatment groups), ^++++^*p* < 0.0001 (reflecting the difference in object exploration) based on post-hoc test following repeated measures ANOVA, revealing a significant main effect of object (*F*_(1,40)_ = 7.45, *p* < 0.01) and an object × treatment interaction (*F*_(1,40)_ = 25.66, *p* < 0.0001). Scatter plot: \*\*\*\**p* < 0.0001 based on unpaired two-tailed *t*-test (*t*_(40)_ = 5.2). **(D)** Absolute exploration times of the conspecific (mouse) and inanimate dummy object (dummy) (line plot) and the social interaction index (scatter plot) in the SI. Line plot: \*\**p* < 0.01 (reflecting the difference between treatment groups), ^++++^*p* < 0.0001 (reflecting the difference in object exploration) based on post-hoc test following repeated measures ANOVA, revealing a significant main effect of mouse (*F*_(1,40)_ = 114.8, *p* < 0.0001) and a mouse × treatment interaction (*F*_(1,40)_ = 12.05, *p* < 0.01). Scatter plot: \*\**p* < 0.01 based on unpaired two-tailed *t*-test (*t*_(40)_ = 3.05). **(E)** Schematic representation of the experimental design used for adult manipulations. Animals received bilateral injections at P84, daily VEH or CNO treatment from P93 to P113, and were behaviorally tested at > P136. *N* = 10 animals per group. **(F)** Total distance moved in the entire arena (left) and center zone (right) during the OF. **(G)** Absolute exploration times of the temporally remote and recent objects (line plots) and temporal order memory index (scatter plot) in the TOMT. ^+^*p* < 0.05 based on repeated

Temporal order memory, a function critically dependent on the mPFC^60,61^, was then assessed. Mice were hereby first allowed to explore a pair of identical objects (sample phase 1), followed by exposure to a new pair of identical objects (sample phase 2). In the subsequent test phase, mice were presented with one object from the first sample phase (temporally remote) and one from the second sample phase (temporally recent). Mice with an intact mPFC were expected to discriminate between the two objects, spending more time exploring the temporally remote object^60,61^. Consistent with this expectation, VEH-treated mice preferentially explored the temporally remote object, exhibiting a positive temporal order memory index (**Fig. 2C**). In contrast, CNO-treated mice failed to discriminate between temporally remote and recent objects **(Fig. 2C)**, indicating persistent deficits in temporal memory following adolescent astrocyte stimulation.

Social behavior, another PFC-dependent function^62^, was evaluated using a modified three-chamber interaction test^63–66^. Although both groups preferred the conspecific over a dummy object, CNO-treated mice spent significantly less time interacting with the stranger mouse, resulting in a reduced social interaction index (**Fig. 2D**), consistent with impaired sociability.

Additional control experiments confirmed that the behavioral deficits were specific to hM3DGq-mediated astrocyte activation and not resulting from CNO treatment *per se*. Indeed, C57BL/6 mice receiving CNO without hM3DGq expression showed no changes in locomotion, temporal memory, or social behavior (**Supplementary Fig. S3A-D**). Importantly, identical astrocyte stimulation applied during adulthood (P93-P113) (**Fig. 2E**) or later developmental periods (P50-P70) (**Supplementary Fig. 3E**) did not produce any deficits in these behavioral and cognitive domains (**Fig. 2F-H; Supplementary Fig. S3F-H**), identifying adolescence as a critical window of vulnerability during which transient overactivation of prefrontal astrocytes is sufficient to disrupt adult PFC-dependent cognitive and social functions.

### Transient overactivation of prefrontal astrocytes during adolescence reduces PV expression and disrupts fast-spiking interneuron activity during gamma oscillations in the adult mPFC

To explore the mechanisms underlying these behavioral deficits, we focused on PV interneurons, which are known to critically regulate PFC-dependent cognitive functions^67^. PV interneurons were first assessed using immunohistochemistry for PV expression and for perineuronal nets (PNNs) (**Fig. 3A**). PNNs are functional extracellular matrix structures surrounding these metabolically active cells that contribute to synaptic stabilization, regulation of excitability, and restriction of plasticity^68,69^. Immunohistochemical analyses revealed a selective reduction in PV intensity in CNO-treated mice irrespective of PNN presence (**Fig. 3B**), whereas PV cell number (**Fig. 3C,E**) and gross morphology (**Supplementary Fig. S4A-B**) were unchanged. PNNs surrounding PV neurons were unaffected (**Fig. 3D**), suggesting that PV dysregulation occurs independently of extracellular matrix alterations. Control mice receiving CNO alone showed no PV changes, confirming the specificity of hM3DGq-mediated astrocyte activation (**Supplementary Fig. S4C-D**).

**Figure 3.**
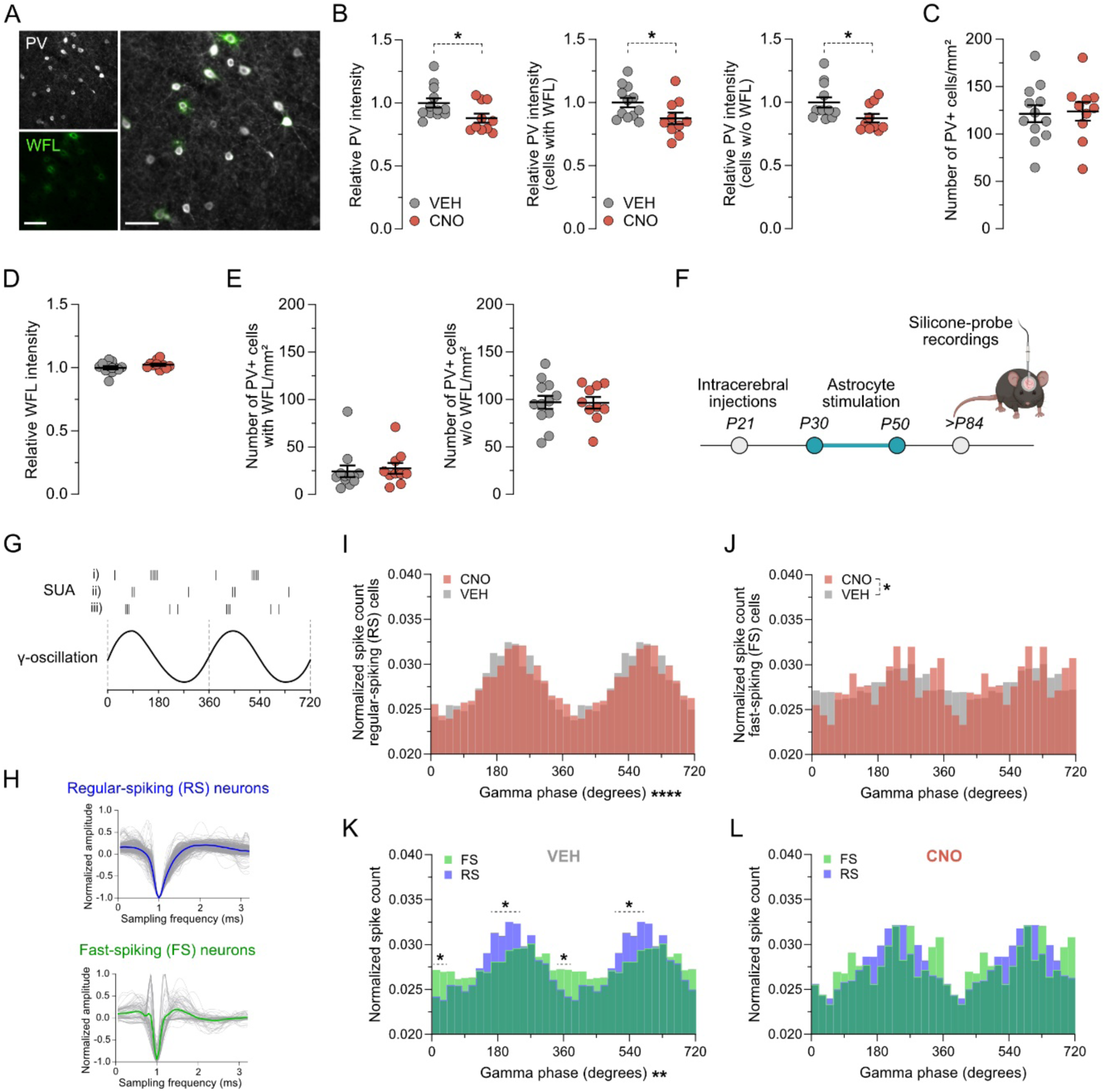
Adolescent prefrontal astrocyte stimulation reduces PV expression and disrupts fast-spiking interneuron activity during gamma oscillations in the adult mPFC. **(A)** PNNs were visualized using wisteria floribunda lectin (WFL). Representative wide-field image of PV^+^ (white) and WFL^+^ (green) cells acquired in the PrL of adult ADAM mice. Scale bar = 50 µm. **(B)** Relative PV intensity (mean gray values, MGV) measured in all PV-expressing cells, as well as in PV-expressing cells with and without (w/o) surrounding perineuronal nets (PNN, WFL^+^). \**p* < 0.05 based on unpaired two-tailed *t*-test (*t*_(20)_ = 2.29). **(C)** Number of PV-positive cells per mm^2^. **(D)** Relative WFL intensity (MGV) surrounding PV-positive cells. **(E)** Number of PV-positive cells with and without (w/o) surrounding PNN (WFL^+^) per mm^2^. For B-E all scatter plots show individual mice (N = 10-12 animals per group) with group means ± SEM. **(F)** Schematic of experimental setup for *in vivo* electrophysiological recordings in urethane-anesthetized adult ADAM mice. A silicon probe was implanted into the mPFC, which recorded neuronal activity from 16 sites for 30 minutes at 20’000 Hz. **(G)** Simplified scheme of single unit activity (SUA) analyses, whereby the normalized spike counts were assessed along gamma oscillation. **(H)** Detected spikes were clustered using wavelet-based feature representations of their waveforms into regular-spiking (RS, *n* = 417) and fast-spiking (FS, right, *n* = 89) cells. **(I)** Group means of normalized spike counts of RS-cells during gamma oscillation. For the purpose of illustration, gamma phase and corresponding spike counts are depicted twice. \*\*\*\**p* < 0.0001 based on two-way ANOVA with repeated measures, revealing a significant main effect of time in gamma (*F*_(17,102)_ = 8.116). **(J)** Group means of normalized spike counts of FS-cells during gamma oscillation. For the purpose of illustration, gamma phase and corresponding spike counts are depicted twice. \**p* < 0.05, based on two-way ANOVA with repeated measures, revealing a significant main effect of treatment (*F*_(1,6)_ = 12). **(K)** Group means of normalized spike counts of RS- and FS-cells during gamma oscillations in VEH-treated mice. For the purpose of illustration, gamma phase and corresponding spike counts are depicted twice. \*\**p* < 0.01 based on two-way ANOVA with repeated measures, revealing a significant main effect of time in gamma (*F*_(17,51)_ = 2.55, p < 0.01). \**p* < 0.05 based on Tukey’s post-hoc test for multiple comparison following significant time in gamma × cell type interaction (*F*_(17,51)_ = 2.71, p < 0.01) **(L)** Group means of normalized spike counts of RS- and FS-cells during gamma oscillations in CNO-treated mice. For the purpose of illustration, gamma phase and corresponding spike counts are depicted twice. For H-L, *N* = 4 animals per group.

Given that alterations in PV expression can define functional properties of FS interneurons^70–73^, we assessed the functional consequences of astrocyte stimulation on neuronal network activity using *in vivo* silicon probe recordings in anesthetized adult mice (**Fig. 3F, Supplementary Fig. S4E**). While power spectral analysis across all frequency bands revealed no differences between VEH- and CNO-treated mice (**Supplementary Fig. S4F-G**), we detected significant changes in single-unit activity (SUA) along gamma oscillations (**Fig. 3I-L**). After spike detection, events were clustered using wavelet-based feature representations of their waveforms^74^, enabling clear separation of regular-spiking (RS) and FS cells (**Fig. 3H**). Normalized spike counts within the corresponding gamma phase (**Fig. 3G**) revealed significant alterations in the activity pattern of FS but not RS neurons in CNO-treated mice (**Fig. 3I-J**). Moreover, FS neurons in CNO-treated mice exhibited altered phase-locking relative to RS bursts^75^ (**Fig. 3K-L**), suggesting impaired coordination of FS interneuron activity within the local network. Together, these findings demonstrate that transient overactivation of prefrontal astrocytes during adolescence leads to long-lasting reductions in PV expression and disrupts the timing precision of FS interneuron activity, which are associated with persistent deficits in PFC-dependent temporal memory and social behaviors.

### Overactivation of prefrontal astrocytes during adolescence transiently disrupts excitatory synapse maturation and PV interneuron input

To further investigate synaptic mechanisms underlying the observed deficits induced by ADAM, we examined the impact of adolescent astrocyte activation on prefrontal maturation. To this end, we performed postmortem analyses on brain tissue collected at P40, the midpoint of CNO treatment (**Fig. 4A**). Based on the adolescent increase in PV protein expression^26^ and the reduction in PV levels observed in adult ADAM mice, we first asked whether astrocyte stimulation acutely alters PV expression. PV immunoreactivity and PV cell density were unaltered at P40 (**Fig. 4B-C**), suggesting that the persistent adult PV deficits (**Fig. 3**) arise gradually and are not evident during adolescence.

**Figure 4.**
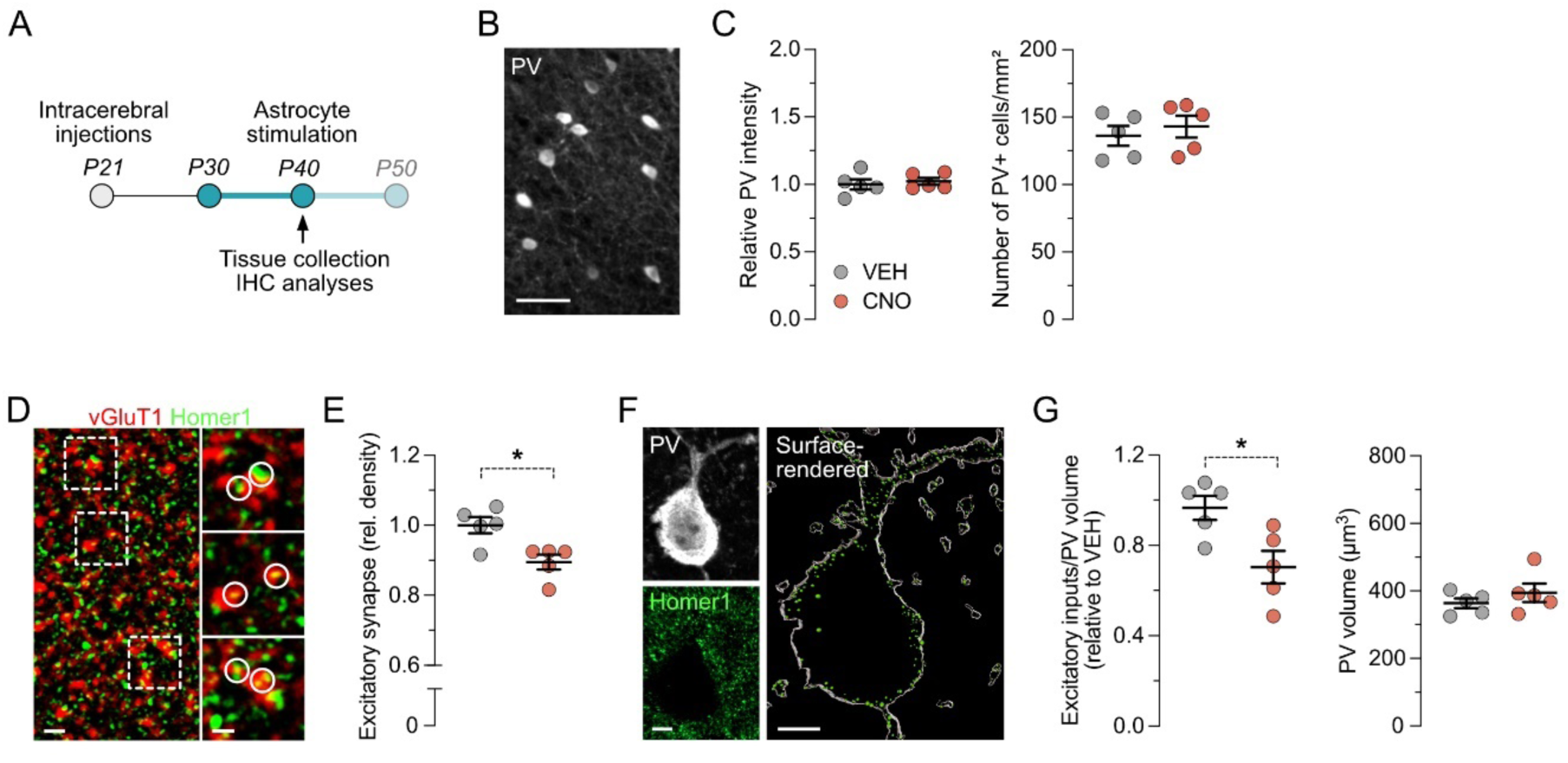
Adolescent astrocyte stimulation transiently disrupts excitatory synapse maturation and PV interneuron input. **(A)** Schematic representation of the experimental design. Animals received bilateral injections at P21 and daily VEH or CNO treatment during adolescence from P30 to P40, upon which postmortem tissue was collected to assess acute effects of ADAM. **(B)** Representative wide-field image of PV^+^ cells acquired in the PrL of P40 ADAM mice. Scale bar = 50 µm. **(C)** Relative PV intensity (mean gray values, MGV) measured in all PV-expressing cells (left) and number of PV-positive cells per mm^2^ (right) counted in mPFC. **(D)** Representative double immunofluorescence stain against vGluT1 (red) and Homer1 (green). vGluT1^+^/Homer1^+^ colocalizations are highlighted with circles and quantified as excitatory synapses. Scale bar overview = 2 µm. Scale bar zooms = 1 µm. **(E)** Scatter plot depicts the relative density of excitatory synapses in VEH- and CNO-treated animals. \**p* < 0.05 based on unpaired two-tailed *t*-test (*t*(8) = 3.34). **(F)** Representative double immunofluorescence stain against PV (white) and Homer1 (green) and the corresponding surface-rendered image used for the colocalization analyses. Scale bars = 5 µm. **(G)** Scatter plot on the left depicts the relative number of excitatory inputs onto PV cells, normalized to the PV cell volume (µm^3^). Scatter plot on the right depicts the average PV volume. \**p* < 0.05 based on unpaired two-tailed *t*-test (*t*(8) = 2.92). All scatter plots show individual mice (*N* = 5 animals per group) with group means ± SEM.

Building on the critical importance of excitatory input for PV interneuron maturation and their functional integration into adult PFC circuits^29^, we next assessed excitatory synapse density. Quantification of colocalized vGlut1^+^/Homer1^+^ puncta revealed a significant reduction in excitatory synapses in the adolescent mPFC following CNO-induced astrocyte stimulation (**Fig. 4D-E**). These effects were not observed in CNO-treated adolescent control mice lacking hM3DGq, indicating that synaptic loss reflects astrocyte-specific activation rather than CNO exposure alone (**Supplementary Fig. S5A-C**).

We then examined whether this synaptic reduction translated to decreased excitatory drive onto PV interneurons. For this purpose, surface-rendered PV^+^ cells were analyzed for co-localized excitatory postsynaptic clusters (**Fig. 4F**). Consistent with global synaptic reductions, excitatory inputs onto PV interneurons were significantly diminished following adolescent astrocyte stimulation (**Fig. 4G**). This effect was absent in CNO-treated controls (**Supplementary Fig. S5D**). Notably, neither excitatory synapse density nor PV innervation remained altered in adulthood, indicating that astrocyte-induced synaptic reductions are transient (**Supplementary Fig. S5E-G**).

### Overactivation of prefrontal astrocytes during adolescence selectively enhances microglia-mediated phagocytosis

Postnatal developmental synaptic refinement is known to be orchestrated by both astrocytes^39,40,76^ and microglia^42,43,77^. We therefore asked whether transient synaptic loss is modulated via glia-dependent synaptic phagocytosis. First, we assessed direct astrocyte-mediated engulfment by quantifying excitatory postsynaptic density positive/lysosome-associated membrane protein 2 positive (PSD-95^+^/Lamp2^+^) puncta within surface-rendered S100β^+^ astrocytes (**Fig. 5A; Supplementary Fig. S6A**). No significant change was observed between control and manipulated animals (**Fig. 5B-D**), indicating that astrocytes themselves do not increase phagocytic removal of excitatory synapses in response to hM3DGq activation.

**Figure 5.**
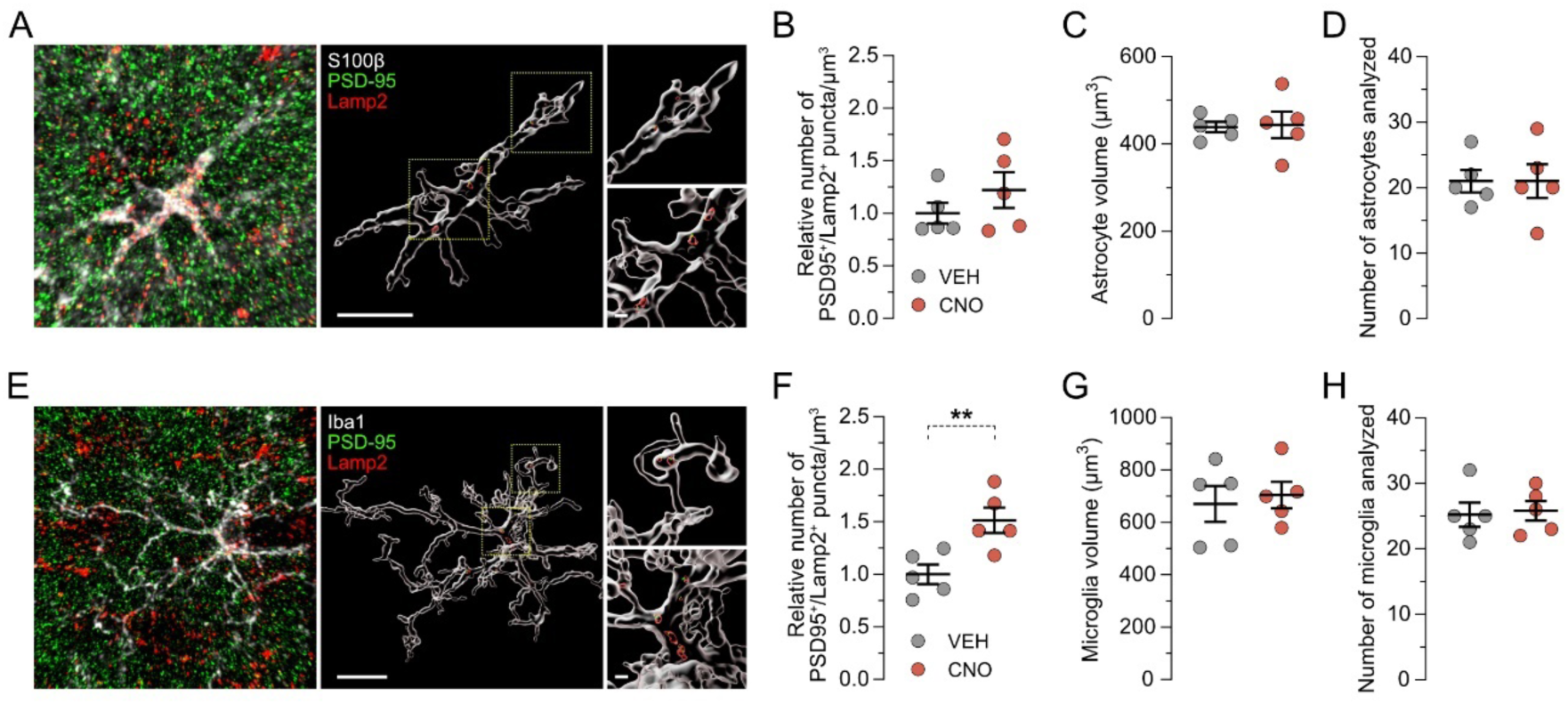
Adolescent astrocyte stimulation increases microglia-mediated synaptic phagocytosis. **(A)** The photomicrographs show a representative triple-immunofluorescence image against S100β (astrocytes, white), PSD-95 (synaptic particles, green), and Lamp2 (lysosomes, red) before (left) and after (bottom) surface rendering and reconstruction. Scale bars = 10 µm and 1 µm. **(B)** Relative number of PSD-95^+^/Lamp2^+^ puncta localized within astrocytes per µm^3^ volume of astrocytes. **(C)** Average volume (µm^3^) of analyzed astrocytes. **(D)** Number of astrocytes analyzed per animal. **(E)** The photomicrographs show a representative triple-immunofluorescence image against Iba1 (microglia, white), PSD-95 (synaptic particles, green), and Lamp2 (lysosomes, red) before (left) and after (bottom) surface rendering and reconstruction. Scale bars = 10 µm and 1 µm. **(F)** Relative number of PSD-95^+^/Lamp2^+^ puncta localized within microglia per volume (µm^3^) of analyzed microglia. \*\**p* < 0.01 based on unpaired two-tailed *t*-test (*t*(8) = 3.37). **(G)** Average volume (µm^3^) of analyzed microglia. **(H)** Number of microglia analyzed per animal. All scatter plots show individual mice (*N* = 5 animals per group) with group means ± SEM.

Because astrocytes can regulate developmental microglia-dependent pruning^41^, we next analyzed PSD-95^+^/Lamp2^+^ puncta within surface-rendered Iba1^+^ microglia (**Fig. 5E; Supplementary Fig. S6B**). In contrast to astrocytes, adolescent CNO treatment significantly increased microglial engulfment of excitatory synapses (**Fig. 5F**), consistent with enhanced microglia-dependent synaptic elimination. This effect was absent in CNO-treated controls lacking hM3DGq, confirming that enhanced microglial pruning is astrocyte-dependent (**Supplementary Fig. S7**).

Collectively, these results demonstrate that transient adolescent astrocyte activation selectively promotes microglia-mediated synaptic pruning, leading to a temporary reduction in excitatory synapse density and PV interneuron input. The transient nature of these effects suggests that astrocytes modulate short-term synaptic remodeling, fine-tuning circuit maturation without permanently altering prefrontal synaptic architecture.

## Discussion

Our study provides causal evidence that astrocytes actively contribute to the protracted maturation of the mPFC. Using ADAM, our adolescent DREADD-based astrocyte model, we show that transient overactivation of prefrontal astrocytes during a defined window of adolescence (P30-P50) is sufficient to induce long-lasting deficits in FS PV interneurons and PFC-dependent cognitive functions in adulthood. Longitudinal Ca^2+^ imaging in awake mice confirmed that repeated oral CNO administration during adolescence significantly increased astrocytic Ca^2+^ while recordings in adulthood (P>84) showed no persistent alterations in astrocytic Ca^2+^ dynamics, indicating that ADAM enables a transient astrocyte stimulation without enduring changes in adult astrocyte signaling. While we cannot exclude the possibility that repeated hM3DGq-mediated astrocyte stimulation induces lasting, Ca^2+^-independent changes in astrocytic functions^78^, such changes are unlikely to account for the cognitive and behavioral deficits observed here. This is supported by the finding that the identical activation paradigm during later developmental stages - early adulthood (P50-70) and adulthood (P91-P113) - failed to elicit behavioral impairments. Thus, our findings identify adolescence as a window of heightened vulnerability during which aberrant astrocyte activity can produce long-term functional consequences.

The specificity of adolescent astrocyte manipulation suggests interference with ongoing maturational processes. Adolescent mPFC development is characterized by remodeling of the inhibitory GABAergic system^26–31^. Key processes include an increase in PV expression^26,79^ and the formation of PNNs around fast spiking interneurons^68,80^. These developmental changes coincide with the emergence of gamma-frequency oscillations^32–34^, which are critical for higher-order cognitive processes^81–83^. Recent evidence suggests that astrocytes play a key role in regulating critical period closure and PV interneuron maturation^44^, raising the possibility that adolescent astrocyte activation impairs cognitive development by disrupting PV interneuron maturation and integration into adult circuits. Consistent with this notion, ADAM induced long-lasting deficits in PV interneurons during adulthood, as reflected by reduced PV expression and altered FS neuron activity within the gamma rhythm, evidenced by impaired phase-locked spiking after RS neuron bursts^75^.

Despite these cellular deficits, we did not detect overt alterations in average mPFC network activity, including gamma-band power, in anesthetized adult ADAM mice. This finding was unexpected, given that the generation and modulation of cortical gamma oscillations depend on PV interneurons^84,85^. However, all recordings were conducted under urethane anesthesia, a condition known to profoundly alter cortical network states, including neuronal excitability and oscillatory dynamics^86–88^. Such state-dependent effects of anesthesia may have masked subtle or behaviorally relevant alterations in network activity that depend on active engagement of prefrontal circuits. Future studies employing recordings in awake, task-engaged animals will be necessary to determine how adolescent astrocyte perturbation impacts mPFC network dynamics during cognitive processing.

Notably, astrocyte-induced long-term deficits in PV interneurons occurred without detectable changes in PNNs, suggesting that PV dysregulation is mediated through mechanisms other than astrocyte-dependent extracellular matrix stabilization^44^. One plausible mechanism involves astrocyte-induced alterations in synaptic maturation, resulting in perturbations in synaptic inputs and interneuron activity. Supporting this hypothesis, we revealed a transient reduction in excitatory synapses, including inputs onto PV interneurons, during adolescent astrocyte stimulation. These observations align with prior work in the developing somatosensory system, demonstrating that synapse elimination critically depends on astrocytic Gq-Ca^2+^ signaling^39^. Moreover, our results indicate that in the PFC, astrocytes influence excitatory synapse density indirectly through microglia-dependent phagocytosis. Signaling pathways mediating prefrontal astrocyte-microglia crosstalk may include astrocyte-derived IL-33^41,89^ or complement-mediated tagging of synapses via astrocyte-derived complement C3^38,77,90^. Alternatively, repeated astrocytic Gq activation may induce structural remodeling of perisynaptic astrocytic processes, thereby increasing synapse accessibility and facilitating microglia-dependent pruning^35^.

We acknowledge several limitations of our study. First, all experiments were conducted exclusively in male mice. Future work will be necessary to determine whether our findings generalize to females, particularly in light of the documented sex differences in astrocyte biology^91^, PV interneuron maturation^92^, and adolescent synaptic remodeling^93^. Second, the precise molecular mechanisms linking astrocyte activity to microglial function and PV interneuron maturation remain unresolved. Elucidating the causal pathways connecting these cellular processes to circuit-level dynamics and behavior will be an important focus of future studies.

Notwithstanding these limitations, our study demonstrates that astrocytes actively contribute to adolescent PFC maturation. Importantly, ADAM provides a unique tool to probe vulnerable developmental periods relevant to psychiatric disorders, particularly those that emerge during adolescence or early adulthood^12^. Although dysfunctional astrocytes have been implicated to contribute to behavioral and cognitive impairments associated with psychiatric disorders via synaptic and neuronal dysregulation^94^, our study provides the first evidence that transient alterations in astrocyte activity during adolescence can induce persistent cognitive and behavioral dysfunctions with translational relevance^81,95,96^. Collectively, our findings emphasize astrocytes as active contributors to prefrontal development and position them as integrative hubs capable of sensing environmental cues through diverse G-protein-coupled receptors^54^, guiding prefrontal maturation in both health and disease.

## Materials and Methods

### Animals

Wildtype C57BL6/N male mice (Charles Rivers, Sulzfeld, Germany) were used in all experiments. All animals were housed in groups of 2-5 per individually ventilated cage in a temperature- and humidity-controlled (21°C ± 3°C, 50 ± 10%) specific pathogen-free holding room with controlled lighting conditions. The animals had *ad libitum* access to standard rodent chow food (Kliba 3336, Kaiseraugst, Switzerland) and water. All procedures were approved in advance by the Cantonal Veterinarian’s Office of Zurich. Every effort was made to minimize the number of animals and their suffering. The number of animals used in each experiment is detailed in the figure and supplementary figure legends.

### Timed-breeding procedure

A timed mating procedure was used to determine the exact birth date of the offspring, as previously described^64,97^. Female mice were paired with male mice in a 2:1 ratio. After vaginal plug verification (gestation day (GD) 0), two pregnant females were housed together for the remainder of the pregnancy and until weaning of the offspring (postnatal day (P) 21), with minimal interference to reduce stress.

### Longitudinal *in vivo* Ca^2+^ imaging

#### Microprism and cranial window implantation

To access the prefrontal cortex for *in vivo* two-photon imaging, a right-handed microprism was implanted in the longitudinal fissure of P21 mice using previously established and validated methods^58,98^. After anesthetizing the animal with a triple anesthesia comprising midazolam (5 mg/kg), fentanyl (0.05 mg/kg), and medetomidine (0.5 mg/kg), the animal’s head was shaved, vitamin A (Bausch & Lomb Swiss AG) cream was applied, and the animal was positioned in a stereotaxic frame. Local anesthesia (50 μl of a mixture containing 1 mL lidocaine (10 mg/mL) and 1 mL bupivacaine (5 mg/mL) in 2 mL saline) was applied under the skin. The skin above the midline was carefully cut and removed. The exposed bone was cleaned and prepared for attaching the headplate using a bonding agent (Prime & Bond). A round headplate was then fixed to the skull using light-cured dental cement (Tetric EvoFlow), exposing the skull above the mPFC for craniotomy. The bridging veins in each animal were carefully examined to prevent bleeding. The bone over the mPFC was thinned with a dental drill (rotate, H-4-002), and smaller bone fragments were gradually removed with a bent needle, ensuring the dura remained intact. After completing the craniotomy, a unilateral injection of a 1:1 mixture of AAV9-hGFAP-hM3DGq-mCherry and AAV9-hGFAP-GCaMP6s virus was performed, using an injection volume of 400 nL at an injection rate of 10 nL/s. The microprism (1.5 mm side length and 1 mm width, S-BSL7 Optosigma) was bonded to a circular glass window (3 mm diameter coverslip) with optical adhesive (Norland #81) and then implanted. A small incision was made along the side of the sinus, into which the microprism was carefully inserted. The microprism was slid into the fissure so that its surface pressed against the previously injected hemisphere. The tissue beneath the microprism was compressed but remained intact. Dental cement was used to secure the glass window in place.

#### Awake in vivo two-photon imaging

Animals were trained for awake *in vivo* two-photon imaging after full recovery from surgery and until they were calm under the microscope. Imaging was performed with a custom-built two-photon laser-scanning microscope^99^ using a 16x water-immersed objective (W Plan-Apochromat 16×/1.0 DIC VIS-IR, Zeiss). GCaMP6s and mCherry were excited at 940 nm and 1100 nm using a Ti:sapphire laser (Mai Tai; Spectra-Physics) with power settings between 10 and 30 mW. Fluorescence emission was detected with a GaAsP photomultiplier module (Hamamatsu Photonics) fitted with either a 520/550 nm bandpass filter or a 607/670 nm bandpass filter, separated by a 560 nm dichroic mirror (BrightLine; Semrock). The two-photon laser-scanning microscope was operated with a customized version of “ScanImage” (r3.8.1; Janelia Research Campus). All imaging was conducted on head-fixed awake mice positioned on an air-lifted platform. High-resolution images (512 x 512 pixels) were captured at 0.74 frames per second, with 20-frame averages for each spot at the beginning of each imaging session to verify consistent localization over multiple weeks. For Ca^2+^ response recordings, images (512 x 512 pixels) were acquired at 1.48 frames per second without averaging. Imaging was performed at baseline (10 minutes) and after administration of VEH or CNO (15 minutes) via MDA. To administer VEH or CNO, imaging was briefly paused and immediately resumed after administration (∼30 s between administration and imaging start). Animals were treated daily with their respective treatment (VEH or CNO) and imaged once per week during the treatment period, as well as once in adulthood (10-minute recording without treatment).

#### Analysis of two-photon images

Whole-frame Ca^2+^ signal analysis was performed using ImageJ. The GCaMP6f signal was normalized to the hM3DGq signal for each frame. For each session, baseline F was set as the average pixel intensity during the session’s baseline activity (900 frames, 10 min). This baseline value was then subtracted from each pixel intensity to calculate ΔF/F for each frame. For event-based analysis of the adult recording, GCaMP6s image sequences were denoised using the “Unsupervised Deep Video Denoising” (UDVD) method^100^, trained directly on the raw images. For training and inference, the image sequences were split into substacks of 128×128×9 (x, y, t) and normalized with z-scores. After denoising, motion correction was applied to all sequences. Subsequently, ROIs were manually assigned to randomly selected astrocytes within the field of view. Calcium events within each ROI were detected using the event-based analysis tool AQuA (version 1.0)^59^, implemented in MATLAB (MathWorks, 2024b). Each recording was manually checked to ensure accurate event detection, and the signal detection threshold was adjusted based on differences in noise levels. For each detected event, the maximum ΔF/F value, the total event count per ROI, and the curve duration were extracted and averaged per ROI.

### Bilateral intracerebral injection

Stereotaxic surgery was performed as previously described^42,43^. All animals received buprenorphine analgesia (0.1 mg/kg s.c.) 30 minutes before surgery. The animals were induced with 5% isoflurane (ZDG9623V, Baxter, Switzerland) in oxygen and maintained under anesthesia at about 1-2% isoflurane throughout the procedure. After shaving the animal’s head and applying vitamin A (Bausch & Lomb Swiss AG) cream, the animals were placed in a stereotaxic frame. Local anesthesia (50 μl of a mixture containing 1 mL lidocaine (10 mg/mL) and 1 mL bupivacaine (5 mg/mL) in 2 mL saline) was applied under the skin. A small incision was made to expose the skull, which was cleaned of connective tissue. A small hole was drilled above the target area using a micro drill (OmniDrill35, World Precision Instruments, USA) with a rose burr (ø 0.3 mm). Mice received bilateral injections into the mPFC based on established coordinates for their age: P21 (anteroposterior [AP] = +1.6 mm, mediolateral [ML] = +0.3 mm, dorsoventral [DV] = −1.35 mm) P41 and P84 (AP = +1.8 mm, ML = +0.3 mm, DV = −1.5 mm). P21 mice received 300 nl, P41 mice received 400 nl, and P84 mice received 500 nl, all injected at a rate of 5 nl/sec. The needle was held in place for 5 minutes after injection, with a small drop of histoacryl (B. Braun, Switzerland) applied to prevent reflux. Incisions were sutured with a surgical thread (G0932078, B. Braun, Switzerland), and the animals received additional analgesia with carprofen (10 mg/kg).

### Daily treatment to activate hM3DGq

To activate hM3DGq, animals received a daily dose of 1 mg/kg clozapine-N-oxide (CNO, BML-NS105-0025, Enzo Life Sciences, Switzerland) dissolved in 0.9% NaCl (B. Braun, Switzerland). The 1 mg/kg dose was selected based on previous chemogenetic studies in rodents^101–103^. CNO and vehicle (VEH, 0.9% NaCl) were administered using the stress-free micropipette-guided drug administration (MDA) method, which is described in detail elsewhere^55,56^. In awake two-photon imaging experiments, CNO or VEH was given via MDA during the imaging session. After 21 days of daily treatments, the mice were left undisturbed until adulthood (> P84), or, for the adult control animals, until the same time period had passed (34 days).

### Behavioral testing

After reaching adulthood, mice underwent behavioral testing. Tests were spaced by at least 48 hours (inter-test interval), except for the open field and the temporal order memory test, in which the open field served as a habituation for the memory test, with a 24-hour interval. All tests were conducted during the animal’s active phase (dark phase of the inverted light-dark cycle) in standardized rooms provided at the Laboratory of Urs Meyer (Institute of Veterinary Pharmacology and Toxicology at the University of Zurich). The equipment was carefully cleaned with water between each animal trial. Special care was taken to ensure that the animals were calm and well-handled before the behavioral experiments began.

### Open field test

A standard open field (OF) exploration test was used to evaluate spontaneous locomotor activity and innate anxiety-like behavior^104^. The OF arenas consisted of identical boxes (40 x 40 cm, with 35 cm-high walls) made of white polyvinyl chloride (OCB Systems Ltd., UK). A digital camera mounted above the arena captured images at a rate of 5 Hz. The room was kept in dim lighting (approximately 30 lux in the center of the arena). The EthoVision tracking system (Noldus Technology, The Netherlands) was used to monitor the animals’ movement and position within the arena (central zone of 10 x 10 cm). The animals freely explored the arena for 30 minutes before being returned to their home cage. Measurements included the total distance moved in the entire arena and within the central zone.

### Temporal order memory test

The temporal order memory test for objects was used to evaluate the animals’ ability to distinguish the relative recency of different stimuli^60,61^. Objects were placed in the OF apparatus described above. The objects used were counterbalanced across the experiments and groups. Three consecutive 10-minute phases were conducted, each separated by 60 minutes in a resting cage where the animals had free access to food and water. The room was kept dimly lit (approximately 30 lux at the center of the arena) during testing, and the resting cages were kept in darkness.

#### Phase 1

A pair of identical objects (blue aluminum hairspray bottles, 250 ml, 20 cm high) was placed in opposite corners of the OF arena, approximately 5 cm from the walls. The animals were gently placed in the center of the OF arena and allowed to explore the objects for 10 minutes. They were then gently removed from the arena and kept in the resting cage for 60 minutes before starting phase 2.

#### Phase 2

A novel pair of identical objects (LEGO Duplo brick pile, 15 cm high) was positioned in the same spots as the first pair from phase 1. The animals were again gently placed in the center of the arena and allowed to explore for 10 minutes. After removing the animals from the arena, they were kept in the same resting cages as before for another 60 minutes.

#### Test phase

In the test phase, one object from each of the two phases (one hairspray bottle and one LEGO brick pile) was allocated in the same location as in phases 1 and 2. The placement (left/right) of the two objects was counterbalanced across all groups. To begin the test, the animals were gently placed in the center of the arena and allowed to freely explore both objects (remote/recent) for 10 minutes. Afterward, the animals were returned to their home cage.

For each animal, the time spent exploring each object was recorded, and the temporal order memory index was calculated using the formula: [(time spent with phase 1 object)/(total exploration time)] − 0.5. The temporal memory index indicates an animal’s ability to discriminate the relative recency of stimuli, with values greater than 0 indicating correct discrimination. Animals were excluded if they did not explore the objects (in total 3 animals, specified in the respective figure legend).

### Social interaction test

The social interaction test was used to measure social interaction with an unfamiliar interaction partner^62^. The apparatus used was a modified Y-maze made from Plexiglass, consisting of three identical arms (50 cm length x 9 cm width and a height of 10 cm) spaced 120° from each other^63–66^. The maze was elevated approximately 90 cm and positioned in a dimly lit room (approximately 30 lux). All the transparent arms of the Y-maze were covered from the outside with white paper sheets. A camera mounted above the Y-maze was used to capture images at a rate of 5 Hz. Two of the three arms contained rectangular wire grid cages (13 x 8 x 10 cm) with metal bars horizontally and vertically spaced 9 mm apart. In one of the metal cages, an unfamiliar mouse (wildtype C57BL6/N of similar age) was placed, while the other wire cage contained an inanimate black object (“dummy”), resembling a black mouse. To start the test, the animal was gently placed into the third cage-free arm and allowed to explore all three arms for 5 minutes. The placement of the social interaction partner (unfamiliar mouse) or dummy object in the two arms (left/right) was counterbalanced across all groups. Social interaction time was counted as the time the animal spent with their nose in contact with the metal cage. The social interaction index was calculated using the formula: [(time spent with mouse)/(total exploration time of mouse and dummy)] − 0.5. A value greater than 0 indicates a preference for the social interaction with the unfamiliar mouse rather than the dummy object. Animals were excluded if they did not explore the mouse or dummy object (in total 1 animal, specified in the respective figure legend).

### *In vivo* silicon probe recordings

#### Data acquisition and preprocessing

To measure brain activity with multielectrode recordings, a craniotomy was performed. All animals received buprenorphine analgesia (0.1 mg/kg s.c.) 30 minutes before surgery. The animals were induced with 5% isoflurane (ZDG9623V, Baxter, Switzerland) in oxygen and maintained under anesthesia at about 1-2% isoflurane throughout the procedure. The procedure, except for the lack of a virus injection and microprism implantation, followed the method described in the section “Microprism and cranial window implantation”. 30 minutes prior to completing the craniotomy, urethane (1.2 g/kg) was injected subcutaneously (s.c.) to ensure a smooth transition of anesthesia. The silicon probe (A4×8-5mm-100-200-177, NeuroNexus Technologies, Inc., USA), with 4 shanks and 8 recording sites per shank, was inserted sagittally into the mPFC using a micromanipulator for precise placement. Only shanks that colocalized with the viral injection, confirmed via postmortem immunohistochemistry, were included in further analysis. The probe was coated with Dil (1,1′-dioctade-cyl-3,3,3′3′-tetramethylindocarbocyanine, dissolved in 70% ethanol). The reference electrode was placed in the cerebellum. Recording commenced 30-60 minutes after probe insertion to allow the brain to adapt to the mechanical stress. Raw signals were sampled at 20000 Hz for a total of 30 minutes (36 million samples per recording site).

#### Spectral Feature Extraction

To eliminate power line interference, all signals were notch-filtered at 50 Hz using a second-order IIR notch filter with a bandwidth of 1.43 Hz, implemented in MATLAB R2023b (MathWorks Inc.). Power spectral density (PSD) estimates were computed using Welch’s method (2-second windows, 50% overlap, 20000 Hz sampling rate), with frequency resolution extended to 200 Hz, as implemented in MATLAB’s pwelch function. Spectral power was determined for six canonical bands: delta (1–4 Hz), theta (4–8 Hz), alpha (8–12 Hz), beta (12–30 Hz), gamma (30–80 Hz), and high-frequency oscillations (80–200 Hz). To ensure comparability across the different recording sites, band powers were normalized by the total spectral power, excluding the 48–51 Hz range to prevent contamination from the notch filter. This resulted in six normalized band power features for each recording site. The recording sites were averaged per animal, and a two-way ANOVA with repeated measures (**Supplementary Fig. S4E**) or nested t-test (**Supplementary Fig. S4F**) was performed.

#### Spike Sorting

Extracellular recordings were processed using the Wave_Clus spike sorting algorithm^74^, which combines wavelet decomposition with superparamagnetic clustering to isolate single-unit activity. Raw signals were sampled at 20000 Hz, providing sufficient temporal resolution to capture the fast dynamics of action potentials. Spike detection was performed by thresholding the continuous signal, and only negative deflections were selected, consistent with the expected polarity of extracellular spikes recorded with our electrode configuration. Detected events were aligned to their negative peaks and subjected to wavelet transformation, which extracts features that efficiently represent spike shapes while reducing noise. These features were then clustered using the superparamagnetic clustering method implemented in Wave_Clus, which does not require *a priori* assumptions about the number of units and is robust to variability in spike waveforms. Clusters were visually inspected to confirm unit isolation, and those showing refractory period violations or unstable waveform shapes were excluded from further analysis.

#### Spike Time Extraction

Spike timestamps and cluster identities were extracted from the corresponding spike files. For each recording, the maximum cluster index was used to determine the number of putative single units. Spike times were retained in milliseconds (from the original acquisition system). For later classification of neuronal cell type (regular-spiking vs. fast-spiking), the average spike waveform for each unit was computed across all spikes by averaging raw waveforms provided in the spike file.

#### Local Field Potential Preprocessing and Gamma-Band Isolation

LFP signals were sampled at 20 kHz. To extract the gamma-frequency component, the raw signal was filtered using a 4th-order zero-phase Butterworth band-pass filter (30–80 Hz). The filter was implemented using a second-order section representation (filtfilt in MATLAB) to prevent phase distortions. Instantaneous gamma phase was computed by applying the analytic Hilbert transform to the band-passed signal. The angle of the complex analytic signal was used as the continuous gamma phase time series for spike–phase interpolation.

#### Spike-Shape Clustering and Identification of FS and RS Units

To classify units into fast-spiking (FS) and regular-spiking (RS) types, average waveforms were subjected to shape-based clustering using a previously computed clustering with a kmeans clustering algorithm in Matlab using k=10. Units belonging to cluster ID 10 were classified as FS units, whereas cluster IDs 1 and 7 were classified as RS units, based on the distinct spike-width and trough-to-peak features obtained from the waveform clustering algorithm. For data representation and statistical analysis, RS or FS cells were averaged per animal, respectively. *<u>Spike–Gamma Phase Histogram Computation:</u>* For each spike timestamp, the corresponding gamma phase was obtained by linearly interpolating the instantaneous gamma phase time series at the spike time. Phases were converted to degrees (0–360°), and each dataset was duplicated (0–720°) to produce stable polar histograms over two cycles. Statistical analysis was performed using one dataset (0-360°).

### Postmortem immunofluorescent analysis

#### Sample collection

Animals were perfused transcardially with a rate of 20 ml/min using artificial cerebrospinal fluid (CSF, pH 7.4), followed by a 6-hour post-fixation in 4% phosphate-buffered paraformaldehyde (PFA, Electron Microscopy Science, 19210), and cryoprotected in 30% sucrose^105^. All samples were cut into coronal sections using a sliding microtome, yielding a series of 8 sections at 30 μm thickness. Slices were stored at −20°C in a cryoprotective solution (50 mM sodium phosphate buffer (pH 7.4) containing 15% glucose and 30% ethylene glycol; Sigma-Aldrich, Switzerland) until further processing.

#### Immunofluorescence staining

Immunofluorescent staining was conducted following established protocols^105^. In short, brain sections were incubated overnight while constantly shaking (100 rpm) at 4°C with primary antibodies (**Table 1**) diluted in Tris-saline buffer (0.5 M Tris, 1.5 M NaCl) containing 0.2% Triton-X-100 and 2% normal donkey serum (NDS). The next day, the sections were washed three times for 10 minutes in Tris-saline buffer before being incubated in the dark at room temperature with constant shaking (100 rpm) with secondary antibodies (**Table 2**), diluted in Tris-saline buffer with 2% NDS. After incubation, they were washed three more times for 10 minutes in Tris-saline buffer before being mounted on gelatinized glass slides. After drying, a coverslip was placed on the slices using Dako fluorescence mounting media (S3023), and the slides were stored at 4°C until image acquisition.

**Table 1:** List of primary antibodies used for immunofluorescence. Rb, rabbit; ms, mouse; ck, chicken; gp, guinea pig; rt, rat.

| Target | Description | Dilution | Vendor, Cat. No. |
| --- | --- | --- | --- |
| mCherry | rt, monoclonal | 1:1000 | Invitrogen, M11217 |
| S100β | rb, monoclonal | 1:1000 | Abcam, ab52642 |
| NeuN | ms, monoclonal | 1:1000 | Merck Millipore, MAB377 |
| Glial Fibrillary Acidic Protein (GFAP) | ck, polyclonal | 1:2000 | Abcam, ab4674 |
| Parvalbumin (PV) | gp, polyclonal | 1:1000 | Synaptic systems, 195308 |
| Wisteria Floribunda Lectin (WFL) | FITC conjugate | 1:1000 | Invitrogen, L32481 |
| Vesicular glutamate transporter 1 (vGluT1) | gp, polyclonal | 1:4000 | Synaptic systems, 135304 |
| Homer1 | rb, polyclonal | 1:1000 | Synaptic systems, 160003 |
| PSD-95 nanobody | FluoTag488 conjugate | 1:1000 | SYSY antibodies, N3702-At488-L |
| Lamp2 | rt, monoclonal | 1:2000 | Abcam, ab13524 |
| Iba1 | rb, polyclonal | 1:1000 | Wako, 019-19741 |

**Table 2:** Secondary antibodies used for immunofluorescence. Rb, rabbit; ms, mouse; ck, chicken; gp, guinea pig; rt, rat

| Target species (host) | Conjugate | Dilution | Vendor, Cat. No. |
| --- | --- | --- | --- |
| rb (donkey) | Alexa Fluor 488 | 1:1000 | Jackson<br>ImmunoResearch Laboratories,<br>711-545-152 |
| ck (donkey) | Alexa Fluor 488 | 1:1000 | Jackson<br>ImmunoResearch Laboratories,<br>703-545-155 |
| rt (donkey) | Cy3 | 1:500 | Jackson<br>ImmunoResearch Laboratories,<br>712-165-153 |
| ms (donkey) | Alexa Fluor 647 | 1:500 | Jackson<br>ImmunoResearch Laboratories,<br>715-605-151 |
| gp (donkey) | Alexa Fluor 647 | 1:500 | Jackson<br>ImmunoResearch Laboratories,<br>706-605-148 |
| rt (donkey) | Alexa Fluor 647 | 1:500 | Invitrogen, A48272 |
| rb (donkey) | Alexa Fluor 750 | 1:500 | Abcam, ab175731 |

#### Microscopy and immunofluorescent image analysis

To examine hM3DGq construct expression, triple-fluorescent images of S100β, NeuN and hM3DGq-mCherry were captured using the Zeiss laser scanning microscope (LSM) 900 with the oil-immersed 40x objective (NA 1.4, 420762-9700-000, Zeiss, Germany), provided at the Institute of Pharmacology and Toxicology (IPT) at the University of Zurich (UZH). 6 images from 3 consecutive mPFC sections within the area of hM3DGq-expression were acquired at a resolution of 1024 x 1024 (zoom: 0.45, pixel size: 0.35 µm, z-step: 1 µm). Higher-resolution image stacks for representative images were acquired at a resolution of 1024×1024 (zoom: 1.3, pixel size: 0.12 µm, z-step: 0.5 µm). For assessing hM3DGq-construct expression, the number of cells expressing mCherry^+^, S100β^+^, mCherry^+^/S100β^+^, and mCherry^+^/NeuN^+^ was counted within each image using ImageJ. Specificity was calculated by dividing the number of mCherry^+^/S100β^+^ cells by the number of total mCherry^+^ cells. Efficacy was calculated by dividing the number of S100β^+^ cells by the number of mCherry^+^/S100β^+^ cells. Leakage was calculated by dividing the number of NeuN^+^ cells by the total number of mCherry^+^ cells. Averages per animal were calculated and displayed as percentages.

Immunofluorescence images of GFAP were captured at the slide imaging system PhenoImager (formerly Akoya Vetra Polaris) provided by the Center for Microscopy and Image Analysis (ZMB) at UZH. With this system, whole slides could be scanned using the 20x objective (NA 0.86), resulting in a pixel size of 0.4961 µm. All analyses shown are from the whole hM3DGq-expressing region in the mPFC. GFAP levels were measured using ImageJ. For each image, the same threshold was applied, the mean gray value (MGV) was measured, and averages were calculated per animal.

Immunofluorescence images of parvalbumin (PV) and wisteria floribunda lectin (WFL) were captured at the slide imaging system PhenoImager (ZMB, UZH) using the 20x objective (NA 0.86), resulting in a pixel size of 0.4961 µm. All analyses shown are from the whole hM3DGq-expressing region in the mPFC. PV and WFL analyses were conducted using Qupath software (version 05.01, United Kingdom). PV^+^ cell bodies were detected using the in-built cell detection function on the PV channel. For each PV^+^ cell body, an 8 µm expansion was made to identify WFL^+^ cells. PV^+^/WFL^+^ cells were selected based on a WFL threshold within this 8 µm boundary. The detected cell number (with and without WFL) and the MGVs of PV (within the cell bodies) and WFL (within the entire cell boundary) were averaged for each animal.

Immunofluorescence images of vGluT1 and Homer1 were captured using the Zeiss LSM900 (IPT, UZH) with the oil-immersed 63x objective (NA 1.4, 421782-9900-000, Zeiss, Germany). 12 images were captured from 3 consecutive sections of the mPFC. To reduce imaging time and because little variability was found between images, 9 images in 3 consecutive sections were obtained for the CNO control cohort. All images were captured at a resolution of 2048 x 2048 (zoom: 1, pixel size: 0.041 x 0.041 µm). A colocalization analysis of the excitatory synapses was performed using a custom macro (provided by J.M. Fritschy at IPT UZH) as previously described^42,43,64,106^. In short, parameters for Gaussian blur, background subtraction, and a threshold were selected and applied to each channel, ensuring consistency across all images. Colocalizations were identified based on pixel clusters in the postsynaptic channel (Homer1) overlapping with pixel clusters in the presynaptic terminal (vGluT1). Postsynaptic pixel clusters were counted between 0.02 and 1.5 µm. A size cutoff for presynaptic pixel clusters and colocalizations was set at 0.02 µm. The number of colocalizations was then averaged per animal.

Immunofluorescence images of excitatory input (Homer1^+^ puncta) on PV were captured using the Zeiss LSM900 with the oil-immersed 63x objective (NA 1.4, 421782-9900-000, Zeiss, Germany). 9 high-resolution images of PV cells on 3 consecutive mPFC sections were captured at a 2048 x 2048 (zoom: 1.9, pixel size: 0.026 x 0.026 µm, z-step: 0.15 µm). Image analysis of the excitatory input on PV was performed using Imaris (version 10.2.0, Oxford Instruments, United Kingdom), provided by the ZMB UZH. PV cells were reconstructed three-dimensionally using the “surface” creation. Homer1^+^ synaptic puncta were generated using the “spots” creation. Homer1^+^ spots were then filtered to colocalize with PV surfaces using the “shortest distance to” function with a threshold of 0.1 µm. To normalize for the size of the PV cell, the number of colocalizations was divided by the volume of the PV surface per image, and an average number of inputs was calculated for each animal.

Immunofluorescence images for the phagocytosis were captured using the confocal microscope Evident Fluoview 4000 with the oil-immersed 60x objective (NA 1.3, UPLSAPO UPlan S Apo, Evident, Japan), provided at the ZMB UZH. Because the phagocytosis assay requires triple immunofluorescence (Cell type marker, Lamp2, and PSD-95) and hM3DGq-mCherry occupancy in the Cy3 channel, we used the near-infrared spectrum (AF750) for the cell type marker. Additionally, to prevent overlap between host species, we used the PSD-95 nanobody as a presynaptic marker, unlike in the rest of the study, where Homer1 was used. The PSD-95 nanobody was tested for colocalization with the Homer1 antibody (data not shown). 4 to 5 images per animal, capturing at least a total of 20 cells of interest, were taken in hM3DGq-expressing regions of 3 consecutive mPFC brain slices. All images were taken at a resolution of 1024 x 1024 (pixel size: 0.11 x 0.11 µm, z-step: 0.22 µm). Phagocytosis colocalizations were performed using Imaris (version 10.2.0, Oxford Instruments, United Kingdom), provided by ZMB UZH. Cell markers and Lamp2 were reconstructed three-dimensionally using the “surface” creation. PSD-95^+^ synaptic puncta were generated using the “spots” creation. The shortest distance from Lamp2 to the target cell was filtered with a threshold of −0.01 µm to ensure Lamp2 was enclosed within the target cell. PSD-95^+^ spots were filtered based on the closest distance to enclosed Lamp2 (threshold 0). Colocalizations were then normalized to the target cell surface volume per image and averaged per animal. The average cell volume was calculated by dividing the total cell surface volume by the number of cells per image.

In all experiments, the data acquisition and analysis were performed by an experimenter blinded to the experimental groups. All images were taken whenever possible during a single imaging session, with all imaging settings kept constant. However, due to large data volumes, multiple imaging days were often necessary. In this case, all experimental groups were spread across different imaging days to avoid bias. Final illustrations were created in ImageJ or Imaris, with contrast and brightness uniformly adjusted across the entire image. One animal had to be excluded from immunofluorescence processing due to poor perfusion quality.

### Blinding and statistical analysis

All data were collected and analyzed by an experimenter blinded to the experimental condition. All statistical analyses were performed using Prism (version 10.6.1, GraphPad Software, La Jolla, CA, USA). The specific statistical tests used are detailed in the respective figure legends.

## Acknowledgements

We thank all past and present laboratory members for their support in this project. We thank the Viral Vector Facility (VVF) of the Neuroscience Center Zurich (ZNZ), Switzerland, in particular Jean-Charles Paterna and Lazaros Vasilikos, for the production of the rAAVs used in this study. We further thank the Center for Microscopy and Image Analysis (ZMB) at the University of Zurich for their equipment and support throughout the study. This work was financially supported by the Swiss National Science Foundation (grant No. PZ00P3_202149 awarded to T.N.). Additional financial support was provided by the Brain & Behavior Research Foundation (grant No. 30963 awarded to T.N.), the Neuroscience Center Zurich (ZNZ PhD Grant 2022 awarded to T.N), and the Olga-Mayenfisch Foundation (awarded to T.N.).

## Contributions

J.F., V.B., A.C., E.B., S.M.S., A.Ö.A., S.F., A.P., O.K., and F.V.M. were involved in the acquisition, analysis, and interpretation of the data. J.F. and T.N. were involved in the conception and design of the study, as well as in the analysis and interpretation of the data. T.N., B.W., T.K., and L.R. supervised the research. J.F. and T.N. wrote the initial manuscript draft. All authors contributed to the review and editing of the final manuscript and have given their approval for the version to be published.

## Ethics declaration

All authors declare no competing interests. The funding of this study was independent of the study design, data collection, analysis, and the decision to publish the manuscript.

## Data availability

Source data is provided within this manuscript or in the supplementary materials. Any additional data we inadvertently missed will be shared upon reasonable request.

