## Supplementary Information for "Adolescent Astrocyte Dysregulation Impairs Prefrontal Interneuron Maturation and Adult Cognition"

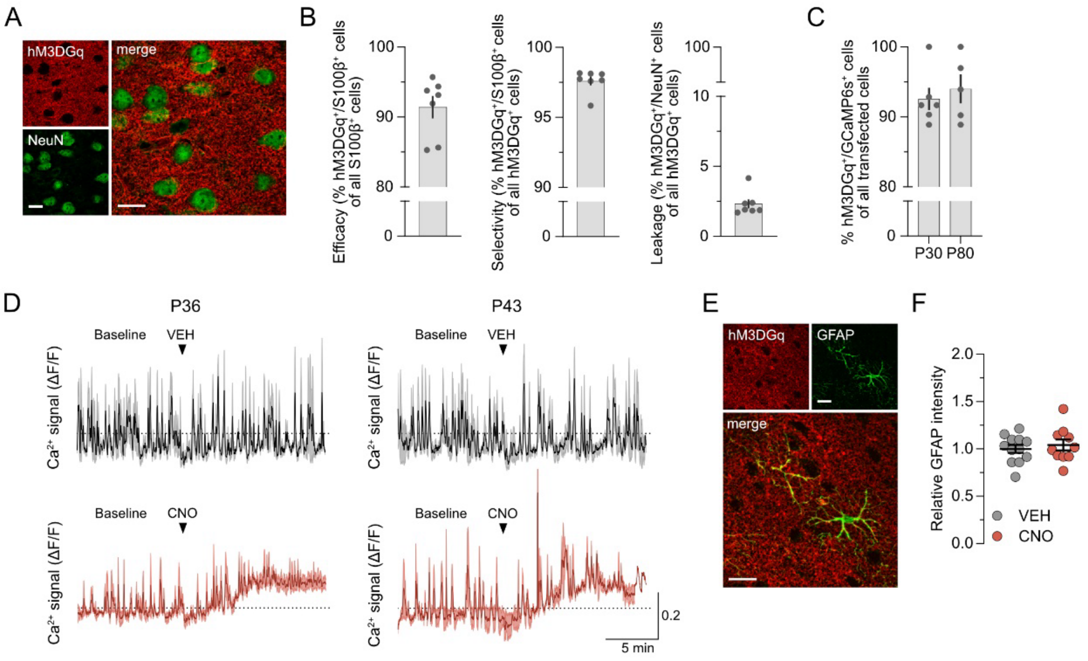


**Supplementary Figure S1. Selectivity and effectiveness of the adolescent DREADD-based astrocyte model (ADAM).**
**(A)** Representative double-immunofluorescence image of neurons (NeuN, green) and hM3DGq (red). Scale bars = 20 µm. **(B)** Scatter bar plots depict efficacy (left), selectivity (middle), and leakage of hM3DGq expression in the adult mPFC. *N* = 7 animals. **(C)** Scatter plot depicts the percentage of cells expressing hM3DGq and GCaMP6s of all transfected astrocytes at P30 and P80. *N* = 6 animals at P30 (*N*(VEH) = 3, *N*(CNO) = 3) and *N* = 5 animals at P80 (*N*(VEH) = 3, *N*(CNO) = 2). **(D)** Whole-frame $Ca^{2+}$ signal (ΔF/F) in VEH and CNO-treated mice at P36 (left) and P43 (right), showing a 10-min baseline followed by a 15-min post-treatment period. *N* = 3 animals per group, except for P43, where *N* = 2 for CNO. **(E)** Representative double-immunofluorescence staining against GFAP (green) and mCherry (red, hM3DGq). Scale bars = 20 µm. **(F)** Relative GFAP intensity (mean grey value, MGV) in the mPFC of adult hM3DGq-expressing mice that received CNO or VEH during adolescence (P30-P50). *N*(VEH) = 12 animals, *N*(CNO) = 10 animals.

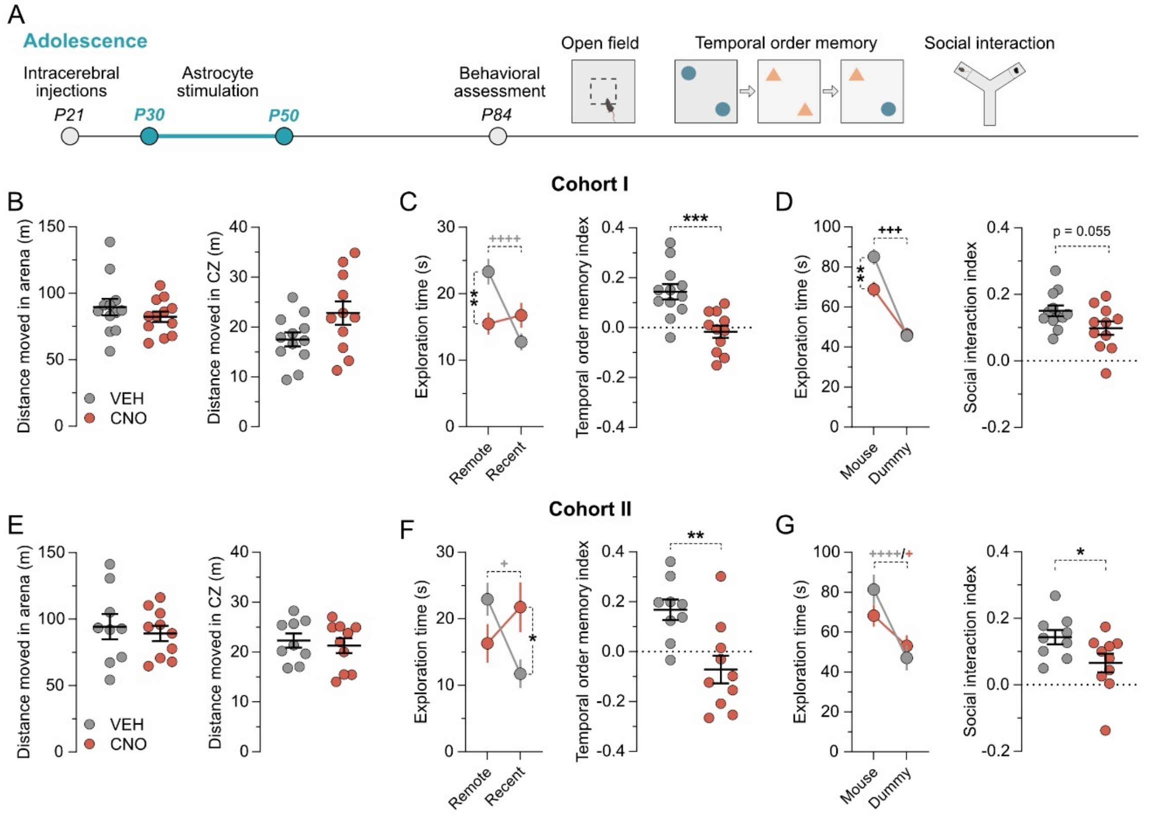


**Supplementary Figure S2**. **Long-term behavioral deficits in two independent ADAM cohorts.** **(A)** Schematic of experimental setup. Animals received bilateral injections at P21, daily VEH or CNO (1 mg/kg, oral via MDA) treatment during adolescence from P30 to P50, and were behaviorally assessed in the open field test (OF), the temporal order memory test (TOMT), and the social interaction test (SI) in adulthood (> P84). Results from Cohort I ($N$(VEH) = 12 and $N$(CNO) = 11) are shown in B-D, and results from Cohort II ($N$(VEH) = 9 and $N$(CNO) = 10) are shown in E-G. **(B)** Total distance moved in the entire arena (left) and center zone (right) during the OF. **(C)** Absolute exploration times of the temporally remote and recent objects (line plots) and temporal order memory index (scatter plot) in the TOMT. Line plot: $^{**}p < 0.01$ (reflecting the difference between treatment groups), $^{++++}p < 0.0001$ (reflecting the difference in object exploration) based on post-hoc test following repeated measures ANOVA, revealing a significant main effect of object ($F_{(1,21)} = 13.21$, $p < 0.01$) and an object × treatment interaction ($F_{(1,21)} = 21.36$, $p < 0.001$). Scatter plot: $^{***}p < 0.001$ based on unpaired two-tailed $t$-test ($t_{(21)} = 4.06$). **(D)** Absolute exploration times of the conspecific (mouse) and inanimate dummy object (dummy) (line plot) and the social interaction index (scatter plot) in the SI. Line plot: $^{**}p < 0.01$ (reflecting the difference between treatment groups), $^{+++}p < 0.001$ (reflecting the difference in object exploration) based on post-hoc test following repeated measures ANOVA, revealing a significant main effect of mouse ($F_{(1,21)} = 87.45$, $p < 0.0001$), significant main effect of treatment ($F_{(1,21)} = 4.71$, $p < 0.05$) and a mouse × treatment interaction ($F_{(1,21)} = 6.53$, $p < 0.05$). Scatter plot: p = 0.055 based on unpaired two-tailed $t$-test ($t_{(21)} = 2.04$). **(E)** Total distance moved in the entire arena (left) and center zone (right) during the OF. **(F)** Absolute exploration times of the temporally remote and recent objects (line plots) and temporal order memory index (scatter plot) in the TOMT. Line plot: $^{*}p < 0.05$ (reflecting the difference between treatment groups), $^{+}p < 0.05$ (reflecting the difference in object exploration) based on post-hoc test following repeated measures ANOVA, revealing a significant object × treatment interaction ($F_{(1,17)} = 9.43$, $p < 0.01$). Scatter plot: $^{**}p < 0.001$ based on unpaired two-tailed $t$-test ($t_{(17)} = 3.42$). **(G)** Absolute exploration times of the conspecific (mouse) and inanimate dummy object (dummy) (line plot) and the social interaction index (scatter plot) in the SI. Line plot: $^{+}p < 0.05$ (CNO-treated mice) and $^{++++}p < 0.0001$ (VEH-treated mice) (reflecting the difference in object exploration) based on post-hoc test following repeated measures ANOVA, revealing a significant main effect of mouse ($F_{(1,17)} = 33.58$, $p < 0.0001$) and a mouse × treatment interaction ($F_{(1,17)} = 5.03$, $p < 0.05$). Scatter plot: *$p < 0.05$ based on unpaired two-tailed $t$-test ($t_{(17)} = 2.16$). All scatter plots show individual mice with group means ± SEM. All line plots show group means ± SEM.

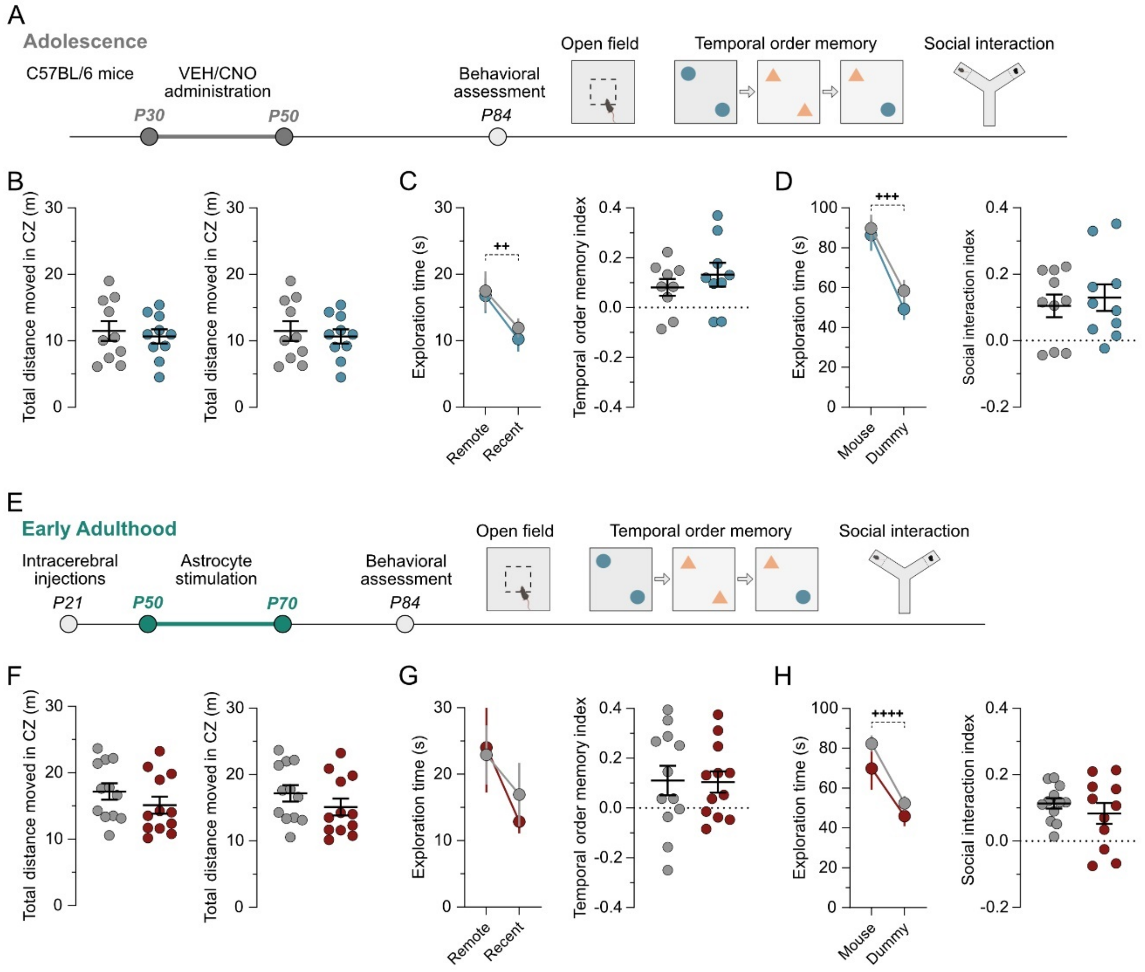


**Supplementary Figure S3**: **CNO-treatment alone and astrocyte stimulation during later developmental periods do not produce long-lasting behavioral deficits.**

**(A)** Schematic of experimental setup for CNO-treatment controls. C57BL/6 male mice received daily VEH or CNO (1 mg/kg, oral via MDA) treatment during adolescence from P30 to P50 and were behaviorally assessed in the open field test (OF), the temporal order memory test (TOMT), and the social interaction test (SI) in adulthood (> P84). *N* = 10 animals per group. **(B)** Total distance moved in the entire arena (left) and center zone (right) during the OF. **(C)** Absolute exploration times of the temporally remote and recent objects (line plots) and temporal order memory index (scatter plot) in the TOMT. Line plot: $^{++}p < 0.01$ based on repeated measures ANOVA revealing a significant main effect of object ($F_{(1,16)} = 9.06$, $p < 0.01$). One animal of each group was excluded because they did not explore the objects. **(D)** Absolute exploration times of the conspecific (mouse) and inanimate dummy object (dummy) (line plot) and the social interaction index (scatter plot) in the SI. Line plot: $^{+++}p < 0.001$ based on repeated measures ANOVA, revealing a significant main effect of mouse ($F_{(1,18)} = 19.62$, $p < 0.001$). **(E)** Schematic of experimental setup. Animals received bilateral injections at P21, daily VEH or CNO treatment from P50 to P70, and were behaviorally tested at > P84. *N* = 12 animals per group. **(F)** Total distance moved in the entire arena (left) and center zone (right) during the OF. **(G)** Absolute exploration times of the temporally remote and recent objects (line plots) and temporal order memory index (scatter plot) in the TOMT. **(H)** Absolute exploration times of the conspecific (mouse) and inanimate dummy object (dummy) (line plot) and the social interaction index (scatter plot) in the SI. Line plot: $^{++++}p < 0.0001$ based on repeated measures ANOVA revealing a significant main effect of object ($F_{(1,21)} = 30.46$, $p < 0.0001$). One animal in the CNO group was excluded because it did not explore the objects. All scatter plots show individual mice with group means ± SEM. All line plots show group means ± SEM.

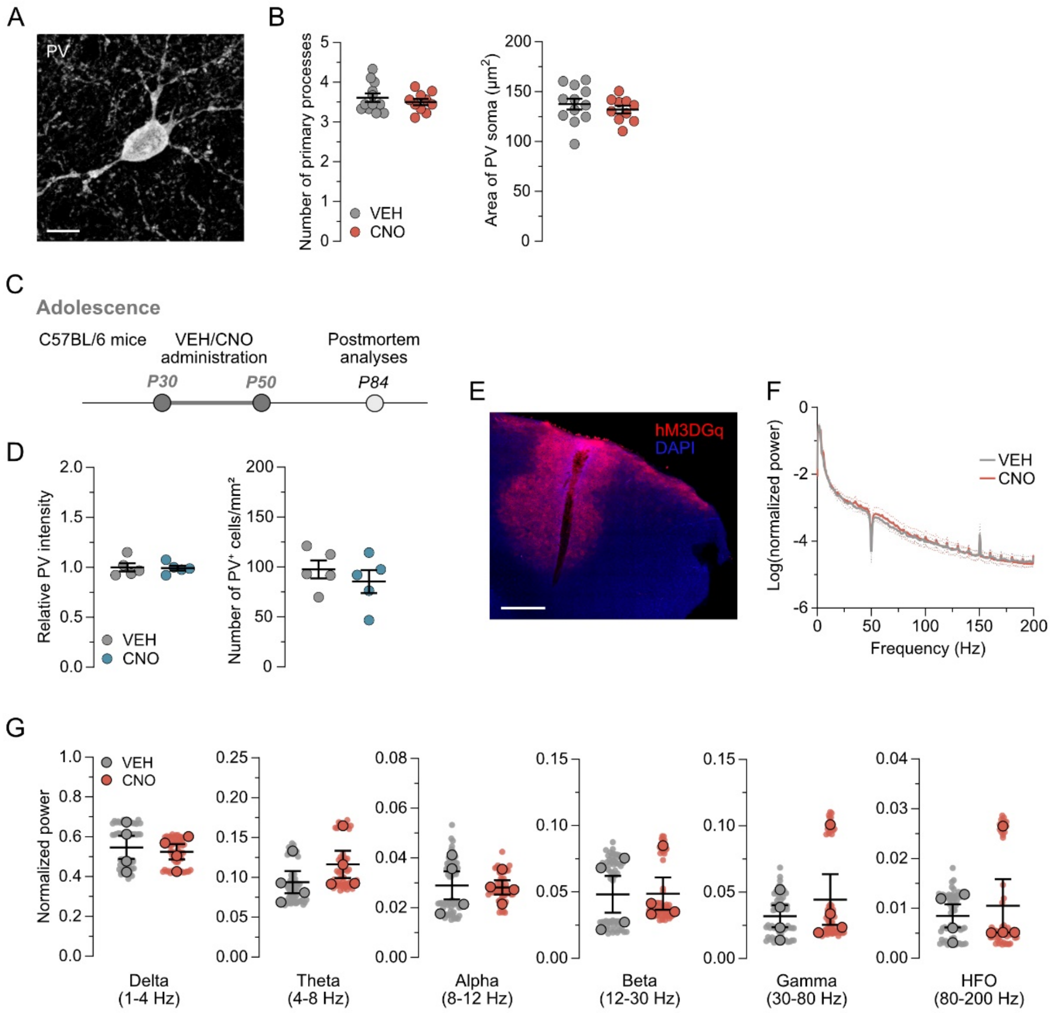


**Supplementary Figure S4**: **Confocal analysis of PV interneurons in adult animals with manipulated astrocytes during adolescence.**

**(A)** Representative high-resolution z-stack image of a PV⁺ interneuron for morphological analyses. Scale bar = 10 μm. **(B)** Scatter bar plots depict the average number of primary processes (left) and the average area of PV soma ($\mu m^2$, right). Data are group means (*N*(VEH) = 12, *N*(CNO) = 10) ± SEM. **(C)** Schematic of experimental setup for CNO-treatment controls. C57BL/6 male mice received daily VEH or CNO (1 mg/kg, oral via MDA) treatment during adolescence from P30 to P50. Tissue for postmortem analyses was collected once the animals reached adulthood (> P84). **(D)** Relative PV intensity (mean grey values, MGV) measured in all PV-expressing cells (left) and number of PV-positive cells per $mm^2$ (right) in the mPFC of adult CNO-control mice. *N* = 5 animals per group. **(E)** Representative image of a multielectrode insertion site in hM3DGq-expressing (red) region of the mPFC (sagittal section, DAPI in blue). Scale bar = 500 μm. **(F)** Log-transformed normalized power spectrum at frequencies between 0 and 200 Hz. The data are group means ± SEM. **(G)** Normalized power in distinct oscillatory bands. Delta (1-4 Hz), theta (4-8 Hz), alpha (8-12 Hz), beta (12-30 Hz), gamma (30-80 Hz), and high-frequency oscillations (HFO, 80-200 Hz). Scatter plots depict the averaged signal over the 30-min recording session per recording site (small dots, *n* = 16 per animal) and overall animal mean (large dots, *N* = 4 animals per group).

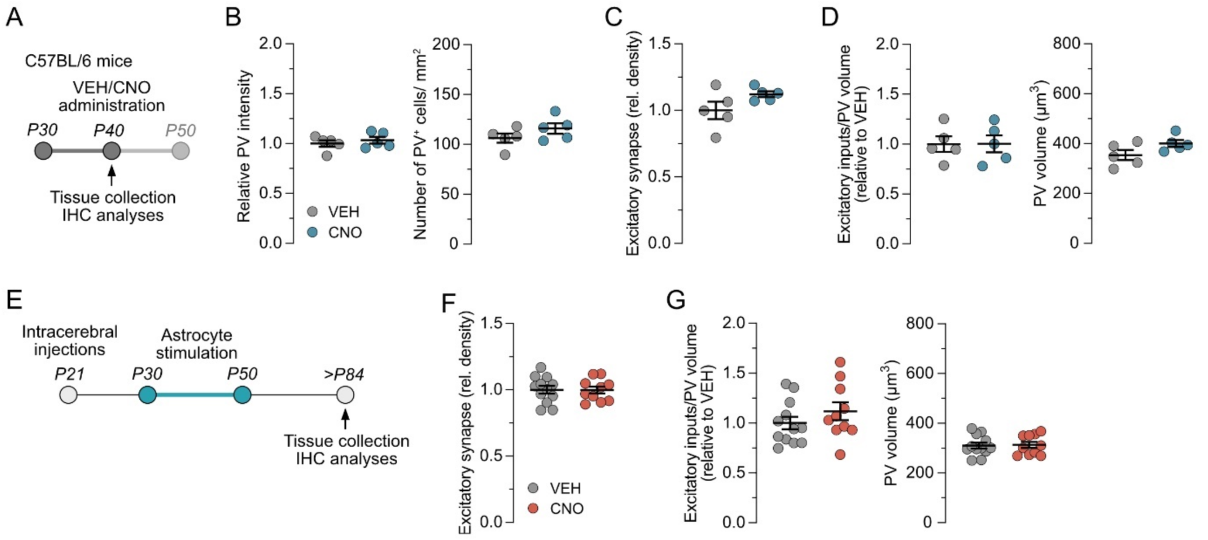


**Supplementary Figure S5: Postmortem analyses assessing acute effects of CNO treatment in control mice and long-term effects of ADAM on excitatory synapses and excitatory input onto PV interneurons.**

**(A)** Schematic of experimental setup for CNO-treatment controls. Post-mortem tissue was collected during CNO treatment (P40, $N$ = 5 per group). **(B)** Relative PV intensity (mean grey values, MGV) measured in all PV-expressing cells (left) and number of PV-positive cells per $mm^2$ (right) in the mPFC of adolescent (P40) CNO-control mice. **(C)** Relative density of excitatory synapses in VEH- and CNO-treated control mice. **(D)** Relative number of excitatory inputs onto PV cells (right) and average PV volume ($\mu m^3$, left). **(E)** Schematic of experimental setup for postmortem analyses in adult ADAM tissue ($N$(VEH) = 12, $N$(CNO) = 10). **(F)** Relative density of excitatory synapses in VEH- and CNO-treated control mice. **(G)** Relative number of excitatory inputs onto PV cells (right) and average PV volume ($\mu m^3$, left). All data show individual mice with group means ± SEM.

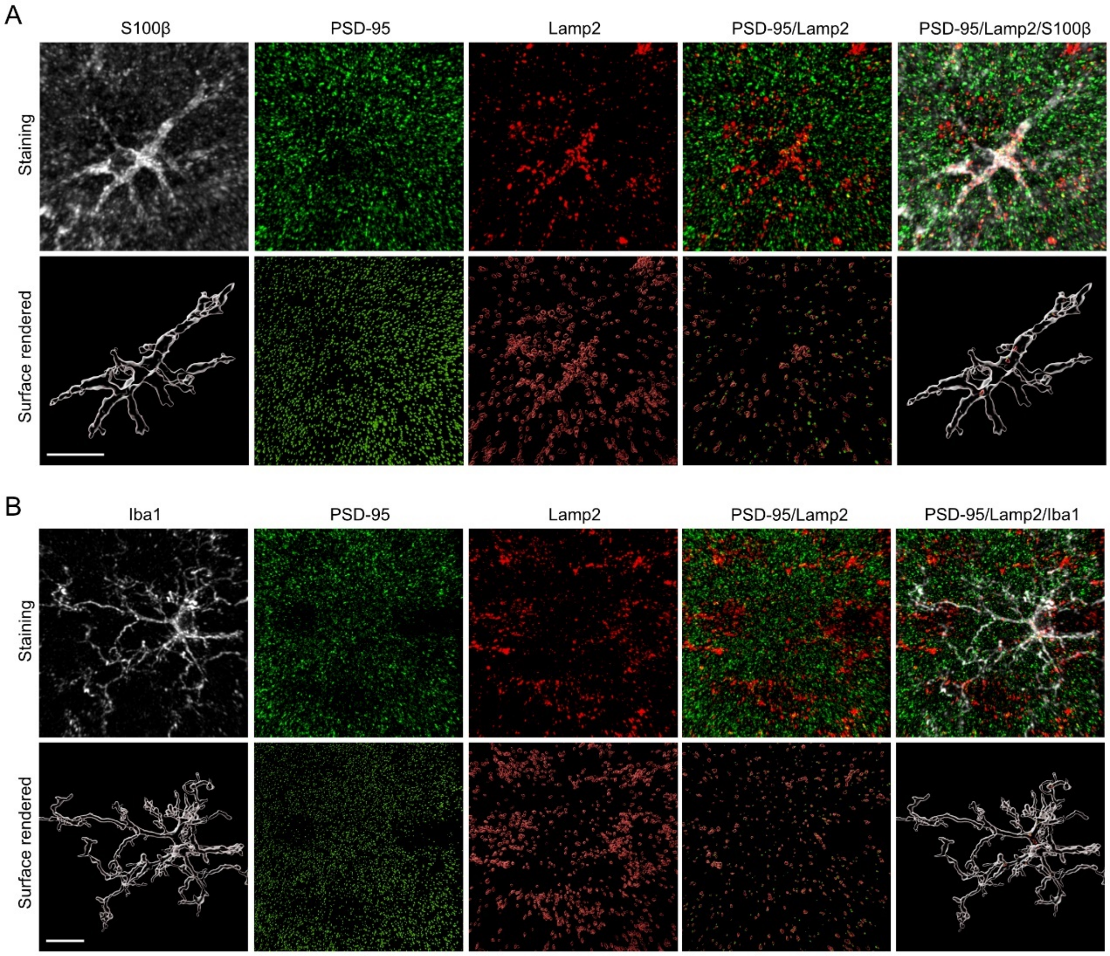


**Supplementary Figure S6: Imaris-based image processing for phagocytic analyses**
**(A)** Representative images used for the Imaris-based colocalization analyses in surface-rendered astrocytes. Upper row depicts IHC stains against the astrocyte marker S100β (white), excitatory postsynaptic density protein PSD-95 (green), and the lysosomal marker Lamp2 (red). Lower row depicts the same images after surface rendering and reconstruction. **(B)** Representative images used for the Imaris-based colocalization analyses in surface-rendered microglia. Upper row depicts IHC stains against the microglia marker Iba1 (white), PSD-95 (green), and Lamp2 (red). Lower row depicts the same images after surface rendering and reconstruction. Scale bar = 10 µm.

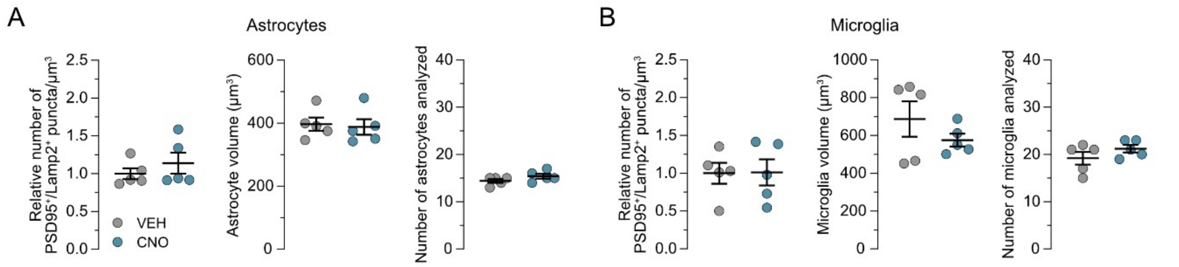


**Supplementary Figure S7: Astrocytic and microglial phagocytosis in adolescent (P40) CNO control animals.**

**(A)** Relative number of PSD-95⁺/Lamp2⁺ puncta localized within astrocytes per µm$^3$ volume of astrocytes (left), average volume (µm$^3$) of analyzed astrocytes (middle), and the number of astrocytes analyzed per animal (right). **(B)** Relative number of PSD-95⁺/Lamp2⁺ puncta localized within microglia per volume (µm$^3$) of analyzed microglia (left), average volume (µm$^3$) of analyzed microglia (middle), and number of microglia analyzed per animal (right). All data show individual mice (*N* = 5 animals per group) with group means ± SEM.
